# No Free Lunch from Deep Learning in Neuroscience: A Case Study through Models of the Entorhinal-Hippocampal Circuit

**DOI:** 10.1101/2022.08.07.503109

**Authors:** Rylan Schaeffer, Mikail Khona, Ila Rani Fiete

## Abstract

Research in Neuroscience, as in many scientific disciplines, is undergoing a renaissance based on deep learning. Unique to Neuroscience, deep learning models can be used not only as a tool but interpreted as models of the brain. The central claims of recent deep learning-based models of brain circuits are that they make novel predictions about neural phenomena or shed light on the fundamental functions being optimized. We show, through the case-study of grid cells in the entorhinal-hippocampal circuit, that one may get neither. We begin by reviewing the principles of grid cell mechanism and function obtained from first-principles modeling efforts, then rigorously examine the claims of deep learning models of grid cells. Using large-scale architectural and hyperparameter sweeps and theory-driven experimentation, we demonstrate that the results of such models may be more strongly driven by particular, non-fundamental, and post-hoc implementation choices than fundamental truths about neural circuits or the loss function(s) they might optimize. We discuss why these models cannot be expected to produce accurate models of the brain without the addition of substantial amounts of inductive bias, an informal No Free Lunch result for Neuroscience. Based on first principles work, we provide hypotheses for what additional loss functions will produce grid cells more robustly. In conclusion, circumspection and transparency, together with biological knowledge, are warranted in building and interpreting deep learning models in Neuroscience.

## 1 Introduction

Over the past decade, deep learning (DL) has underpinned nearly every success story in machine learning, e.g., [57, 6] and increasingly many advances in fundamental science research, e.g., [36]. In neuroscience, DL is similarly gaining widespread adoption as an indispensable method for behavioral and neural data analysis [52, 50, 28, 43, 40, 46].

But DL offers a unique contribution to neuroscience that goes beyond its role in other fields, in that deep networks can be viewed as models of the brain. The success of DL in matching or surpassing human performance suggests it is now possible to construct models of circuits that may underlie human intelligence. As a recent review wrote, “researchers are excited by the possibility that deep neural networks may offer theories of perception, cognition and action for biological brains. This approach has the potential to radically reshape our approach to understanding neural systems” [54]. Further, DL is a democratizing force for building neural circuit models of complex function.

Here, we examine the essential claims (and promises) of DL-based models of the brain, which are that Because the models are trained on a specific optimization problem, if the resulting representations match what has been observed in the brain, then the models reveal which optimization problem(s) the brain is solving, or 2) These models, when trained on sensible optimization problems, should generate novel predictions about the brain’s representations and behavior.

These are extremely valuable potential contributions. However, given the nascent nature of such approaches and the exuberance accompanying some claims in current work, we should examine them carefully. In DL, some successes attributed to novel algorithms have been shown to instead stem from seemingly minor or unstated implementation choices [65, 21, 35]. In this paper, we ask whether Neuroscientists should similarly be cautious that DL-based models of neural circuits that make specific claims about revealing the brain’s optimization functions or that generate specific neural tuning curves may tell us less about fundamental scientific truths and more about programmers’ particular implementation choices, and might be more post hoc than predictive.

To explore these questions, we evaluate recent DL-based models of grid cells in the entorhinal-hippocampal circuit. The medial entorhinal cortex (MEC) and hippocampus (HPC) are part of the hippocampal formation, a brain structure critical for learning and memory. In a pair of Nobel-prize winning discoveries, HPC was shown to contain **place cells** [48], and MEC, its cortical input, was shown to contain **grid cells** [30]. Place cells each fire at one or several seemingly random locations in small and large environments [51], while grid cells fire in a spatially periodic hexagonal lattice pattern in all two-dimensional environments [30]. Over five decades, the hippocampal formation has been central to understanding how the brain organizes spatial and episodic memory, for experimentalists and theorists alike, with many mysteries remaining. A recent series of DL-based models of the circuit [15, 3, 59, 68, 47]) present a story that training neural circuits on the task of **path integration (PI)** (i.e., updating one’s positional estimate by integrating velocity from self-motion), possibly with the addition of a non-negativity constraint on firing rates [59], results in the emergence of grid cells.

We use code from prior publications to demonstrate these results are due not to the core (path integration) task the network was trained to perform but to separate and specific post-hoc implementation choices that implicitly made the known tuning shapes of grid cells part of the target, even though the narrative accompanying many of these papers suggests that the emergence of those tuning curves rather naturally “falls out” from training on the core task. By leveraging theoretically-guided large-scale exploration and hypothesis-driven experimentation, we show:

1. Networks trained on path integration tasks almost always learn to optimally encode position, but almost never learn grid-like representations to do so.
2. The emergence of grid-like representations depends wholly on specifically chosen structural choices of the network and readouts, rather than on the path integration task, and the structural choices are based on the implicit goal of obtaining grid-like responses.
3. Under more-realistic structural choices for the network readouts, grid cells disappear.
4. Even with the structural choices, grid emergence can be hyperparameter and seed sensitive and non-generic.
5. Multiple grid modules, a fundamental characteristic of the grid cell system, do not emerge from path integration.
6. Grid periods and period ratios, contrary to assertions [3], are not determined by the task and are not fundamental properties that can serve as predictions about observed values in the brain; rather, they are set by hyperparameters selected by the programmer.

In short, deep learning models of MEC-HPC yield grid-like units only when a sequence of specific and biologically implausible implementation choices are made to intentionally bake grid-like units into the task-trained networks, and the emergent grid-like units lack key grid cell properties. Given the non-genericity of grid cell emergence in successfully path integrating networks, it is highly improbable that DL models of path integration would have produced grid cells as a novel prediction from task-training, had they not already been known to exist.

Moreover, it is unclear what new understanding the current models contribute, beyond or even up to what has already been shown for these circuits by existing models. Our results challenge the notion that deep networks offer a free lunch for Neuroscience in terms of discovering the brain’s optimization problems or generating novel a priori predictions about single-neuron representations, and warn that caution, transparency, and biological knowledge are needed when building and interpreting such models.

### Code availability

Our work benefited from previous publications’ published code[47]. To facilitate further research, we similarly release ours: github.com/FieteLab/NeurIPS-2022-No-Free-Lunch.

## 2 Background: Grid Cells

Grid cells [30] are found in the medial entorhinal cortex of mammals and are tuned to represent the spatial location of the animal as it traverses 2D space. Each cell fires at every vertex of a triangular lattice that tiles the explored space, regardless of the speed and direction of movement through the space. As a population, grid cells exhibit several striking properties that provide support for a specialized and modular circuit. Grid cells form discrete modules (clusters), such that all cells within a module share a common period and orientation, while different modules express discretely different spatial periods [62]. The period ratios of successive modules have values in the range of 1.2-1.5.

The mechanism underlying grid cells is widely supported to be through attractor dynamics: Translation-invariant lateral connectivity within the grid cell network results in a linear Turing instability and pattern formation [9, 23, 7]. These models explain how grid cells can convert velocity inputs into updated spatial estimates, and make several predictions that have been confirmed in experiments, including most centrally the stability of low-dimensional cell-cell relationships regardless of environment and behavioral state, that define a toroidal attractor dynamics [24, 72, 64, 26, 25].

## 3 Experimental approach

The central messaging of existing DL models of grid cells is that training ANNs on **Path Integration (PI)** – using self-velocity estimates to track one’s spatial position – causes the networks to learn grid cells [15, 3, 59, 47], even when the technical portions and code implementations of the papers involve many other critical choices without which grid cells would not emerge.

Thus, we focus on asking: if a recurrent neural network is trained on PI, what is the approximate probability that it will exhibit grid cells? We follow the setup used by many previous papers: a 2.2 m × 2.2 m arena is created, then, spatial trajectories (i.e. sequences of positions and velocities) are sampled. Networks receive as inputs the initial position and velocities, and are trained to output (some encoding of) the positions in a supervised manner (Fig. 1ab). There are multiple possible encodings of position, and, as we will show, this choice is critical. Two simple encodings are Cartesian [15] or polar [1]. Another encoding scheme is via bump functions in 2D space, with each output assigned different positions that collectively tile the space with similar tuning curve shapes [3, 59, 59, 47]. This encoding has been equated with place cells, even though place cells’ fields tend to be heterogeneous in size and shape [51, 19], as well as in number [51], and, unlike the choice of a difference-of-Gaussians (DoG) or difference-of-Softmaxes (DoS) readout tuning in ANN models, do not exhibit any inhibitory surround. See Appendix A for position encoding details. For all encodings, supervised learning is used to train the network via backpropagation through time.

**Figure 1:**
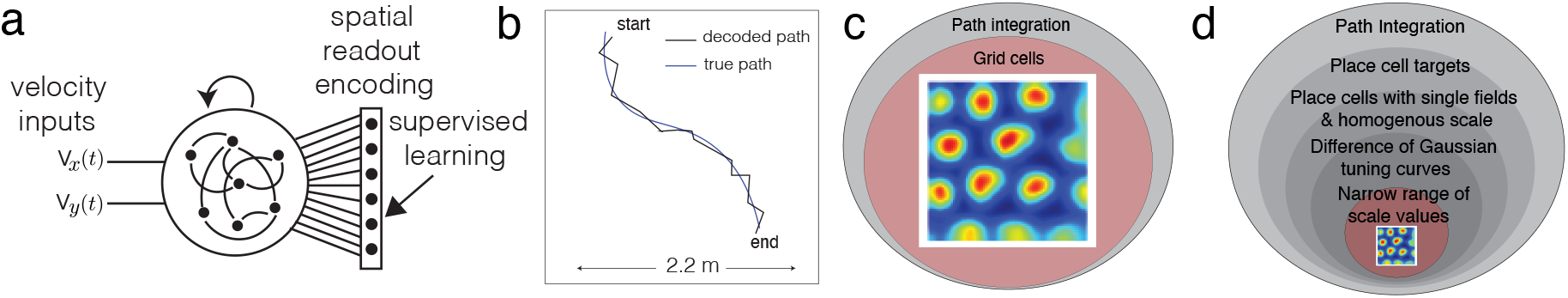
Setup and claim. (a-b) Schematic of recurrent neural network setup to predict some encoding of 2D position from 2D velocity. (b-c) Grid cells in recent DL papers, obtained in part by learning to path integrate [15, 3, 59, 60, 47], conclude that path integration creates grid cells. (d) We show that most artificial neural networks (ANNs) trained to path integrate can do so, but only a very small subset of such networks yield grid cells, and that grid cell emergence results in ANNs are post hoc: they result from post-facto selection of architectures, functions, and hyperparameter settings that specify grid cells as an implicit target.

### Spatial tuning assessments

The spatial tuning ratemaps of hidden units in the networks are the primary basis for comparison with the brain’s grid cells. To compute ratemaps, a trained network is evaluated on long trajectories that cover the 2D environment, then each hidden unit’s average activity per spatial bin is computed. Ratemaps of units are compared with grid cells through the gridness score use by [3, 59, 60, 47]. *We are extremely lenient with classifying a particular network training run a success: if even a single hidden unit has a grid score above a certain threshold, we say the model possibly possesses grid cells*. The grid score, when applied to ANN units without additional criteria, is not perfect since cells classified by grid scores represent only an upper bound on the total number of grid cells (e.g. the high grid score given to units with triangular symmetry without a periodic pattern, Fig. 2d (Gaussian readouts)); for details, see Appendices B and C.

**Figure 2:**
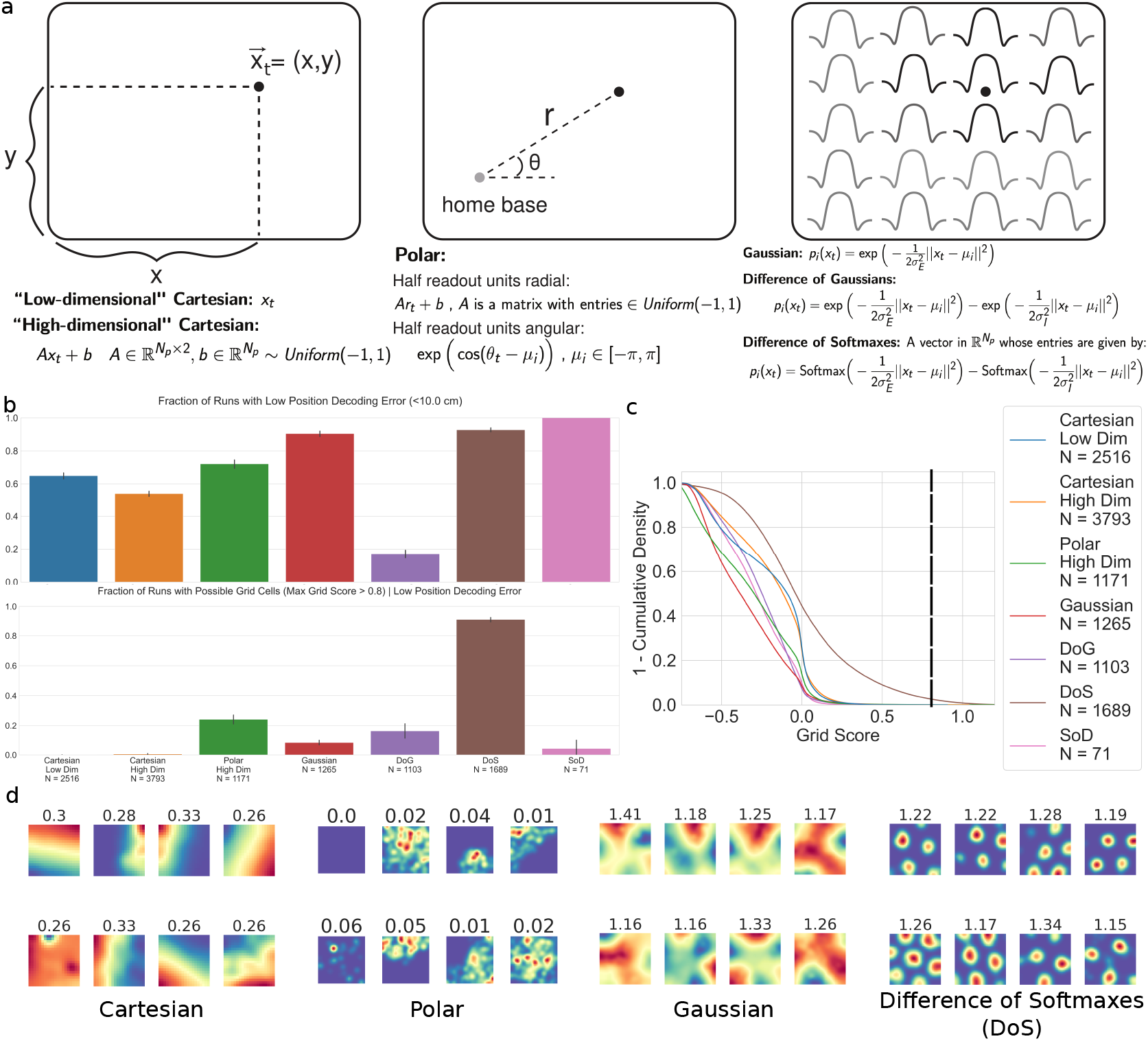
Grid-like response requires highly specific target encoding. (a) Readout encodings of spatial position. (b) Top: Across readout encodings, most networks learn the PI task. Bottom: Few networks display *possible* grid-like representations (threshold = 0.8). (c) Survival functions of grid scores per readout encoding. (d) Rate maps of topgrid-scoring units in ANNs performing good PI with i) Cartesian, ii) Polar, iii) Gaussian, iv) specifically selected (tuned) Difference-of-Softmaxes (DoS) readouts. i)-iii) do not learn grid cells. Numbers above rate maps are 60° grid scores.

## 4 Networks trained on path integration tasks learn to estimate position, but rarely learn grid cells

We demonstrate that most path-integrating networks do not converge to a grid-like solution, instead requiring very specific architectural choices including readout tuning functions. Grid-like representations emerge when the programmer makes choices that, rather than relating to the path integration objective or biologically realistic place cells, are designed post-hoc to produce grid cells.

We ran large-scale hyperparameter sweeps across common implementation choices: 1) Architectures: RNN [20]; LSTM [33]; GRU [13]; UGRNN [14]; 2) Activation: Sigmoid; Tanh; ReLU; Linear; 3) Optimizers: SGD, Adam [41]; RMSProp [32] 4) Supervised Targets: Cartesian; Polar; high-dimensional bump-like readout population code with Gaussian [3], Difference-of-Softmaxes (DoS) [59, 60, 47] or Difference-of-Gaussians (DoG) tuning curves. 5) Loss: mean squared error on the agent’s Cartesian position [37, 15]; geodesic distance on the agent’s polar position [1]; cross entropy on a high-dimensional population of bump-like readout units [3, 59, 47] 6) Miscellaneous: recurrent & readout dropout, initialization, parameter L2 regularization, seed.

For networks with bump-like population readouts, we additionally swept: 1) Width *σ* of Gaussian readouts; 2) Whether the bump-like readouts have homogeneous or heterogeneous field widths; 3) For DoG or DoS readouts, the surround scale s, i.e., the ratio between the inhibitory and excitatory Gaussian standard deviations 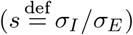 For DoG readouts, the ratio of amplitudes *α*_*E*_/*α*_*I*_ between the two Gaussians. 5) Number of fields per readout unit. Evaluating the entire hyperparameter volume is computationally prohibitive, so we evaluated a subvolume most consistent with previous papers, focusing our exploration around conditions that did produce grid cells. In this sense, our search was biased toward configurations shown to produce grid cell emergence and thus our findings about the fragility of these solutions conservatively favored these solutions as much as possible. All sweeps are provided in Appendix F.

To evaluate whether a network learns to optimally estimate spatial position from velocity inputs, we measured its position decoding error using previous papers’ methods [3, 59, 47]: using the network’s output Cartesian positions (if trained on Cartesian targets) or by decoding position from the network’s outputs. Any network with error < 10 cm was considered to have achieved optimal position encoding.

In total, we trained > 11, 000 networks and found that most succeed in learning to path integrate (Fig. 2a, Top), but few learn grid cell representations (Fig. 2a, Bottom). This is consistent with earlier work [37, 1] demonstrating that networks can learn to path integrate and solve other hard navigational problems (e.g. self-localization across multiple environments and identification of spatial environment from ambiguous cues, a case of self localization and mapping or SLAM) without grid-like units emerging as a solution.

## 5 Grid-like unit emergence requires specific supervised target functions

We next sought to characterize when grid cells are learnt under different encodings of 2D spatial position in the readout units (i.e. supervised targets). We tested multiple encodings: i) Cartesian, ii) Polar, iii) Gaussian, iv) Difference-of-Gaussians (DoG), and v) Difference-of-Softmaxes (DoS).

We found we that in the ANN network architecture of Fig. 1a, grid cells do not emerge from Cartesian or Polar readouts, consistent with earlier work [37]. Similarly, they do not emerge from Gaussian encodings (Fig. 2) [37]. Consistent with this result, the Gaussian readouts of [3] used in tandem with 50% dropout and a different architecture do not yield (square or hexagonal) grid cells without dropout (result not shown). and, as shown recently and independently by [70], although grid-like responses can be obtained with Gaussian readouts after the addition of another constraint, they disappear without. [59, 60] critiqued [3] to show that the hexagonal patterning of cells in [3] was indistinguishable from low-pass filtered noise. However, their [59, 60] focus was to argue that a neural nonlinearity (which they termed a non-negativity constraint, though any non-odd function suffices) robustly produces hexagonal firing – which they showed by replacing the simpler Gaussian-like readouts of [3] with a specific DoG/DoS readout.

We found that DoG/DoS readouts [59, 60] were critical for producing grids (Fig. 2), reproducing the main results of [59], even with non-negativity constraints: Lattices of any geometry (hexagonal or square) only emerge with DoS fields, corresponding to a small and particular subset of DoG fields.

### Grid period values set by hyperparameters, and multiple modules do not emerge

Next, two prominent features of grid cells are their intrinsically set periods (invariant to the external environment) and the existence of a discrete set of grid periods that scale by a rough factor of 1.4 between adjacent scales [62]. Multi-periodicity is critical for unambiguous spatial coding over large scales. We asked whether ANN models generate multiple periods and whether their the period values are fundamental or hyperparameter dependent.

To ensure we would obtain at least some grid cells, we fixed the readouts to be DoS, and swept over different scales *σ*_*E*_ of the DoS. We found that almost all runs had a unimodal distribution of grid periods (Fig. 3a), meaning the networks learnt only one module of grid cells. Contemporaneously with the NeurIPS review process, other researchers independently reported the same result [56].

**Figure 3:**
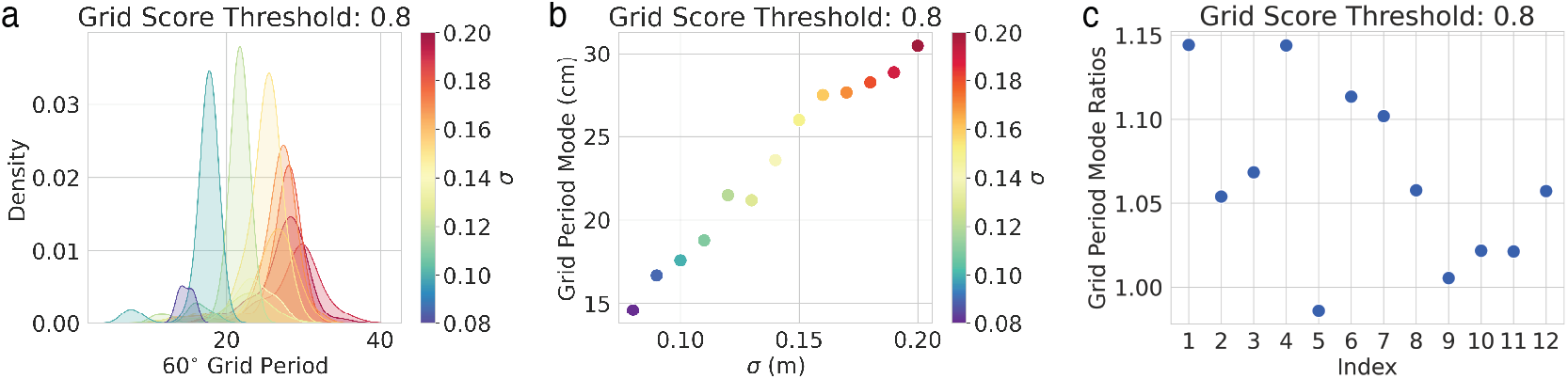
The spatial scale of grids is set by hyperparameters and multiple modules do not emerge. (a) Over a wide sweep of DoS (Fig. 2a bottom) target field widths *σ*_*E*_, the distribution of grid periods is unimodal (each color: period distribution from 3 runs with same *σ*_*E*_ value; different periods are only obtained by varying *σ*_*E*_), meaning multiple grid modules do not emerge, in contrast to the brain’s grid cell circuit. (b) The chosen target field width *σ*_*E*_ determines the grid period mode, meaning that grid period is not a prediction of the models. (c) If we smoothly sweep *σ*_*E*_ as a proxy for simulating different modules, the distribution of adjacent periods produces ratios closer to 1 than the experimental ratios of ∼ 1.4.

Further, we found that the period of formed grid-like representation is completely determined by the width *σ*_*E*_ of the externally imposed readout DoS (Fig. 3) and other specific parameter choices. The period of the grid-like responses increased monotonically with the width of the DoS readout (Fig. 3b). Since individual networks did not learn multiple modules, we used the somewhat discrete distribution of peaks of the single module formed when sweeping the DoS *σ*_*E*_ more continuously to compute grid period ratios. These period ratios from adjacent peaks led to non-biological values (Fig. 3c).

### Other details with DoG/DoS readouts affect grid emergence

In all DL-grid cell papers we examined [3, 15, 59, 60, 47], we discovered implementation details critical to the emergence of grid cells that were not stressed in the claims. As one example, we discovered an implementation detail essential for the emergence of grid cells in a series of papers [59, 60, 47] that is unmentioned in main texts and supplements. These papers report using an unnormalized equi-norm Difference-of-Gaussian (DoG) readout target function ([59] Appendix C1,[60] Methods 4.2, [47] Appendix C1), i.e. a DoG with amplitude parameters set to 1, but their code uses a Difference-of-Softmaxes (DoS) target function. When we trained ideal grid-forming ReLU networks with equi-norm DoG tuning curves, sweeping the receptive field *σ* and surround scale s, they did not result in grids (Fig. 4b). The Fourier analysis (below) explains why equinorm-DoG tuning should not produce lattices (Fig. 4c).

**Figure 4:**
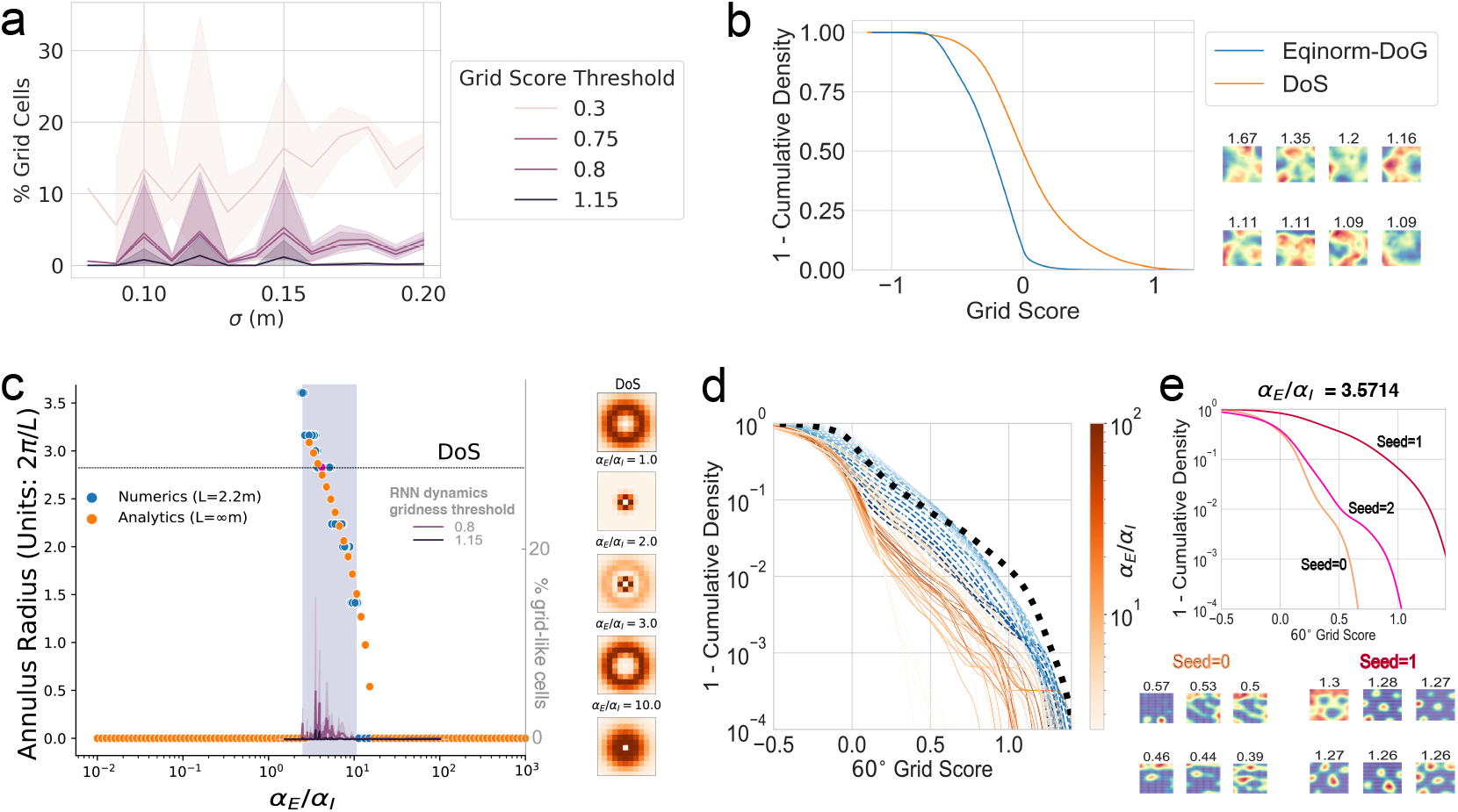
Other details with DoG/DoS readouts affect grid emergence: Even with DoS/DoG translation-invariant targets, grid solutions are sensitive: (a) Modestly varying the DoS width *σ*_*E*_ can cause grids to disappear. (b) Left: Grid scores of networks trained with equinorm-DoG versus DoS readouts [59, 47, 60] shows DoS is critical for high grid scores. Right: Rate maps of top grid scoring units from equinorm-DoG networks. (c) A necessary condition for grid emergence with DoG readouts is that the Fourier transform of the readout correlation matrix contain an annulus of radius > 2π/*L* (*L* is the size of the enclosure in which the RNN is trained). The hyperparameter region where this condition is met is < 1 order of magnitude (FT annulus radius computed analytically (orange) and through numerical construction of fields in finite environment (blue)). RNNs do not learn periodic responses outside this region, but most RNNs inside the region do not either, meaning the annular radius criterion is insufficient. DoS readouts are similar to one particular choice of DoG amplitudes, and only DoG amplitudes very close to the DoS point succeed in producing periodic responses. (d) Across *α*_*E*_/*α*_*I*_ DoG ratios, DoG networks generally score worse than filtered-and-thresholded noise (blue) and worse than DoS (black). (e) More densely sweeping within the theoretically feasible *α*_*E*_/*α*_*I*_ DoG region, and choosing *α*_*E*_ and *α*_*I*_ magnitudes closely matching DoS, shown in (c) still showed only one ratio that did at least as well as a DoS (App. E), but this result was true for one out of three seeds; two “sibling” runs with otherwise identical settings produced poor gridness.

We next trained ReLU RNNs on general DoG readouts, sweeping the component amplitudes *α*_*E*_, *α*_*I*_ while holding *σ*_*E*_, *σ*_*I*_ fixed at ideal values. The theory of [59] predicts that outside the feasible region (blue boxed region of Fig. 4c), no lattices should emerge, but inside the feasible region, all/most RNNs should learn grids. We found the first prediction held, but the second did not: most display grid score distributions comparable to or worse than low-pass-filtered-then-thresholded noise, and well below the ideal-width DoS grid score distribution (Fig. 4d). To investigate, we swept densely inside the feasible region, additionally matching the amplitudes created by DoS (Fig. 4c; Fig. 10); one run out of 1086 surpassed the DoS grid score distribution (*α*_*E*_/*α*_*I*_ *≈* 3.5714, seed=1), but its two “sibling” runs (all hyperparameters same; seeds: 0, 2), Fig. 4e. Thus, and contrary to [59]’s theory based on static function-fitting, DoG readouts at most rarely produce lattices when implemented in actual RNN simulations.

### Fourier analysis of Turing instability provides intuition for the preceding empirical results

Why do only Difference-of-Softmaxes (DoS) or very specific DoG readouts produce grid-like units? We shed light on our present findings by restating the essence of previous analyses of first-principles models [9, 38] here, and leveraging the connection made between these models and trained deep networks in [59]. In the first-principles continuous attractor models, neural dynamics are given by 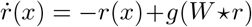, where *x* designates the neural index (in a continuum approximation for neurons), *W* ⋆ *r* designates the total (integrated) inputs from the network to the neuron at index *x*, and *g* is the non-linearity, if the recurrent weight interaction is translationally invariant, then *W* (*x, x*′) = *W* (*x − x*′) = *W* (∆*x*). Under DoG interactions:

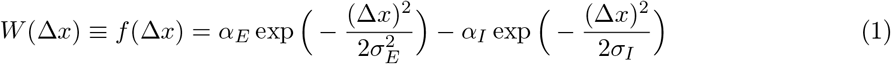

where ∆*x* refers to the difference of indices between the neural pair linked by the weights. The evolution of activity can be decomposed into the growth and decay of Fourier components of the rate vector, which is fully determined by the Fourier transform of *W*, which is given by:

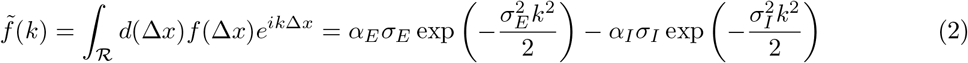

Here *α*_*E*_ (*α*_*I*_) denotes the strength and *σ*_*E*_ (*σ*_*I*_) denotes the scale of excitation (inhibition). For linearized dynamics that approximate 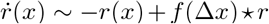 (i.e., *g* has been linearized), the solution will be periodic if the maxima of 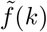, given by 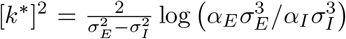, contains sufficient power and if *k*^*^ ≠0. Specifically, the condition for pattern formation is 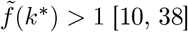 [10, 38]. In particular, the inhibitory surround contained in *f*(∆*x*), with strength *σ*_*I*_, is key to pattern formation; if *σ*_*I*_ → ∞ or *α*_*I*_ → 0, the maximum of the Fourier-transformed weights is at the origin (*k*^*^ = 0), corresponding to a DC (non-patterned/non-periodic) activity state. Similarly, only particular choices of the ratio *α*_*E*_/*α*_*I*_ work, Fig. 4c. In sum, a Gaussian recurrent interaction cannot produce periodic patterns in the continuum limit (relatively large number of cells, large environment), as known from first principles models.

The theory of [59] contributes a connection between these first principles models and feedforward networks performing supervised least-squares regression (“function approximation framework”) onto a target readout *P*, through the observation that gradient optimization of the MSE reconstruction loss ||*P* − *W*_readout_*r* ||^2^ can be approximated as 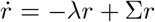, where *σ*_*x,x′*_ = Σ _*i*_ *P*_*i*_(*x*)*P*_*i*_(*x*′)^*T*^ is the specified spatial correlation matrix of the target readouts and *λ* is a regularization parameter. This dynamics now resembles that of first-principles grid cell models, provided the readout correlation matrix has the same form as the first-principles recurrent interaction matrix: *W* (*x, x*′) = *σ*_*xx′*_, upto scaling factors. If the readout target functions are set to be translationally invariant with DoG tuning curves, the readout correlation matrix is a difference of multiple Gaussians (Appendix D), Fig 4c. By the linear stability analysis outlined above, it follows that DoG tuning can sometimes produce grids, but simple Gaussian tuning curves will not generically produce periodic patterns without additional assumptions. This is true whether we consider discretized real and Fourier space or take the continuum limit in both cases: Gaussian readouts generate roughly uniform (non-periodic) activation as the dominant state. Only under the further assumptions of discretization induced by a small spatial environment, higher-order non-linear effects, and orthogonality of hidden units, might Gaussians be theoretically predicted to produce periodic responses [59], and then the periods and shapes of the grids will depend on the environment properties. However, neither hexagonal nor square grids emerge at a significance level beyond filtered and thresholded noise in actual RNNs trained to PI with Gaussian and most DoG readouts.

## 6 Grid cells disappear with realistic readout population heterogeneity

Place cells, to which grid cells project, differ significantly from the highly idealized single-scale, single-field translation-invariant ANN readouts, even ignoring the particular center-surround shape for each field. Place cells have heterogeneous field widths, many with multiple fields [51, 19] and non-uniform spatial correlations. Place cells at similar dorsoventral locations can exhibit a range of field sizes, and single place cells themselves exhibit a diversity of field widths [19]. This naturally leads to the question: Will readout targets with more place cell-like heterogeneous responses still produce grid cells? We found that networks trained with multiple-field, multiple-scale DoS readouts achieve position decoding error as low as single-field single-scale DoS encodings (Fig. 5a), but do not learn grid cells (Fig. 5bc). This finding is consistent with the strong requirement in ANN models of a translation-invariant readout code for grid emergence [3, 59]. Translation invariance is a specific property of grid cells, but it is not likely true of place cells, which as a population over-represent borders, landmarks and reward locations [53, 49, 69, 31, 34, 18, 16, 73, 27, 5]. These observations further demonstrate that ANN models essentially build the known structure of grid cells into their targets, rather than obtaining them from training on simple tasks with plausible readouts.

**Figure 5:**
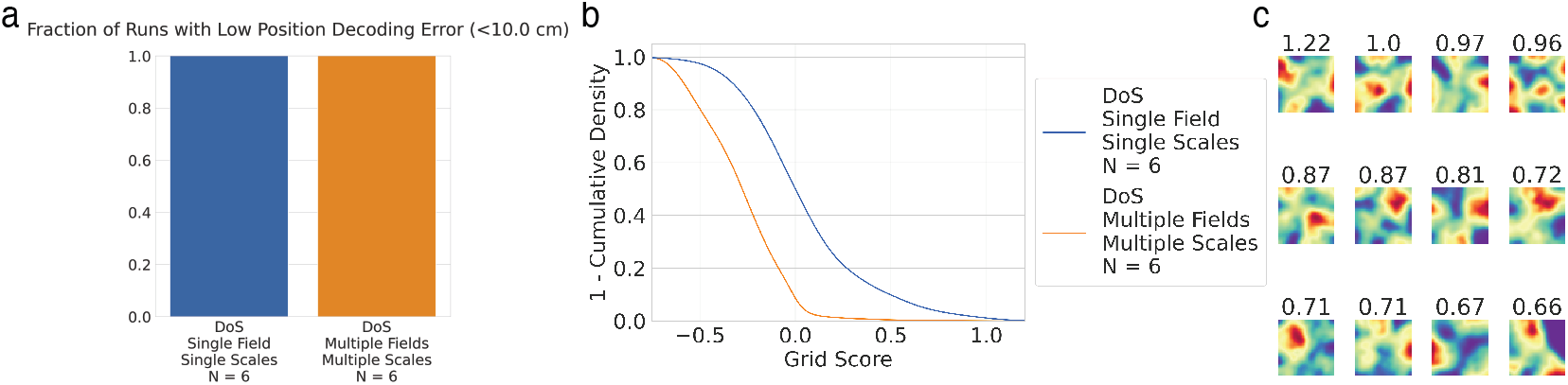
Adding place cell-like heterogeneity to readouts prevents grid emergence. We selected DoS RNNs with the best hyperparameters for grid cell emergence (RNN or UGRNN, ReLU, *σ*_*E*_ = 0.12 cm, *s* = 2.0, 3 seeds), then tested the effect of multiple fields per place cell (∼1 + Pois(3.0)) and multiple scales (receptive field width *σ*_*E*_ ∼ Unif(0.06, 1.0) m and surround scale s ∼Unif(1.25, 4.5)). (a) Networks with multi-scale multi-field DoS readouts all obtain low position decoding error. (b) Multi-scale multi-field DoS readouts do not learn grid cells. (c) Highest-scoring rate maps from multi-field multi-scale networks.

## 7 Why path-integrating ANNs might achieve high predictivity of MEC data

We conclude by introducing a puzzle. A recent NeurIPS spotlight [47] notes that networks trained on single-field single-scale DoS readout encodings explain variance in mouse MEC neural activity at nearly 100% of variance explained by other mice. In contrast, our results demonstrate that these networks learn few grid cells, produce unimodal grid period distributions inconsistent with biological grid cells, and require readout encodings inconsistent with biological place cells to do so. How are these networks able to predict mouse MEC neural activity so well?

The analysis code is not open source, so we are unable to investigate this puzzle. However, we offer a conjecture with preliminary evidence. The analysis of [47] linearly regressed rate maps from one agent (mouse or network) onto rate maps from another mouse, and used Pearson correlation as a measure of “neural predictivity.” We conjecture that different architecture-activation pairs achieve different neural predictivity scores because different pairs learn different intrinsic dimensionalities that then provide richer/poorer bases for linear regressions.

To explore our conjecture, we trained [47]’s 5 architectures (RNN, LSTM, GRU, UGRNN, SRNN) with DoS readouts and ReLU activations (5 seeds per architecture-activation pair). For each trained network, we extracted rate maps and computed a standard linear measure of dimensionality called participation ratio [42], then plotted each pair’s average participation ratio against the published neural predictivity. We found networks with higher dimensional rate maps have higher neural predictivity (Fig. 6).

**Figure 6:**
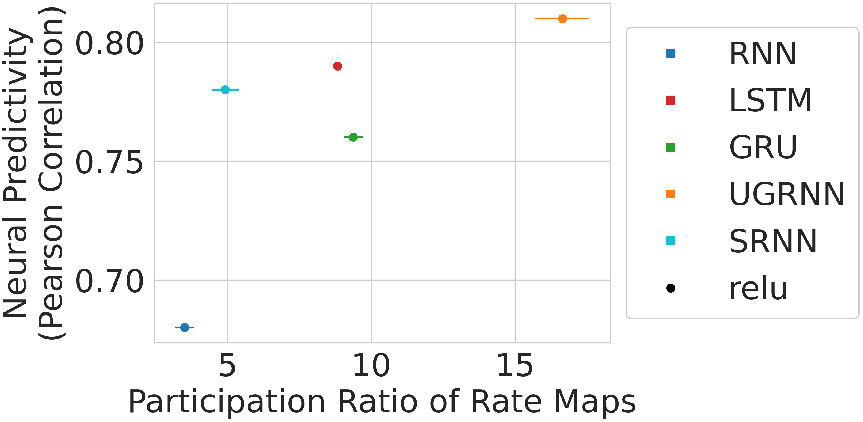
Networks with higher (lower) dimensional rate maps display higher (lower) “neural predictivity” of mouse MEC rate maps. Each dot is an architecture-activation pair; participation ratio averaged over 5 seeds. Neural predictivity from [47].

We caution this correlation between dimensionality and neural predictivity is not (yet) strong evidence. However, we predicted that similar results would be found for other species and modalities, and during the subsequent NeurIPS review process, two independent research groups confirmed our predictions in macaque vision [2] and human audition [66]. If correct, the conjecture raises the interesting research question of whether linear regression-based comparisons of ANNs with neural data might produce better matches to biology more because ANNs have higher dimensional representations than competitors, than because of any detailed similarity.

## 8 Discussion

For research that uses deep networks as models of the brain, there is a fundamental obstacle to making the claim that a given optimization problem is what the brain is solving: If we know the responses of a significant fraction of units from biological networks performing a certain task, we cannot infer the loss function that the brain is optimizing since in principle, numerous different loss functions can have the same/similar minima (Fig. 7 top). In other words, there is typically a *many-to-one* mapping between loss functions and some point in model space where the losses have a minimum. Some of the different grid models from DL and first principles show that this is possible [15, 3, 59, 68, 9]. Conversely, given a reasonable optimization problem that we select based on an organism’s ecological niche, we cannot infer a single solution (and thus build truly predictive single-cell tuning models), since there exist several minima to that optimization problem (Fig. 7 bottom). In other words, there is typically a *one-to-many* mapping from a loss function to its set of solutions.

**Figure 7:**
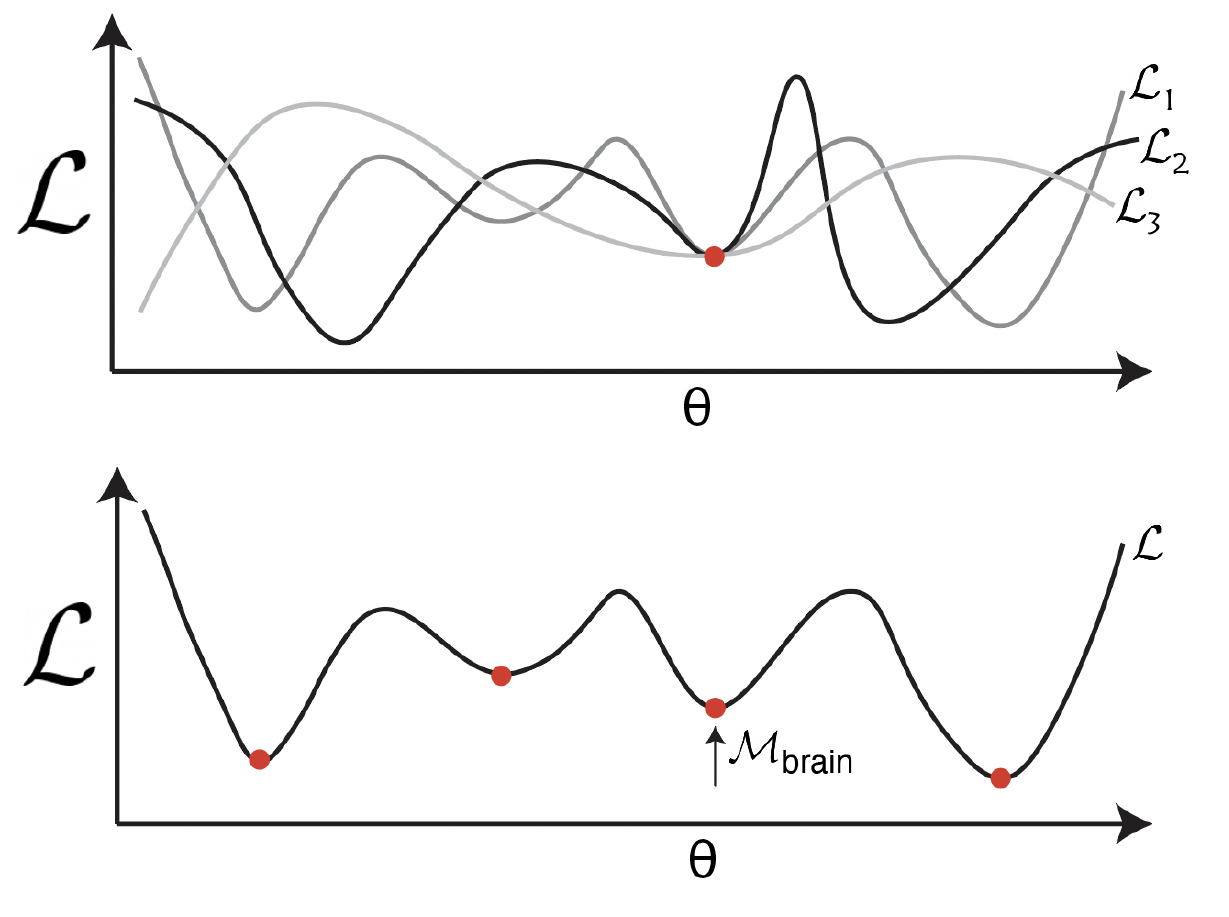
Challenges in achieving the two central claims of recent DL models of neuroscience: Top: Building a model that replicates observed neural responses does not guarantee that the loss function used is the brain’s objective, as multiple objectives can share a solution. Bottom: Training a network on a plausible loss function or even the correct loss function need not yield the solution the brain has selected because the loss function may have multiple minima, of which the brain selects one based on its constraints, while an ANN selects another, based on the optimization technique used.

To break this uniqueness problem and arrive at truly predictive models and a better understanding of the brain’s optimization problems, we must understand the specific inductive biases and constraints present in the biological system we are trying to model. It is untenable to expect success without doing so. This is what we refer to as an informal neural ‘No Free Lunch’.

Can we learn about brain circuits from DL models *without* considering biological inductive biases? Low-dimensional latent representations and dynamics that emerge as necessary for solving difficult problems are possibly sufficiently robust and abstract to be predictive of populations in a neural circuit. For instance, any model solving the task of finding ripe apples in color photos, will create some abstracted representation of round red objects; this would also be a robust prediction for neural systems, but not unique to them. On the other hand, we should generally not expect detailed single neuron tuning correspondences without specific additional constraints or inductive biases: If a low-dimensional latent representation is necessary to solve a task, there are a multiplicity of ways to project it onto the activities of a large number of neurons. Which projection the brain selects depends on factors including energetics, neuron number, downstream uses, and the vagaries of evolutionary dynamics; the projections of ANNs depend on similar factors but specific to the ANN’s loss and the vagaries of gradient descent learning. Consistent with this, models of the visual pathway [71, 4] and circuits that solve latent inference tasks [63, 55, 1, 67] exhibit population-level representations of abstract variables necessary to solve the task. By contrast, DL-based grid cell models make fine-grained claims about single-neuron tuning, which should be surprising without the incorporation of significant additional constraints. Only in cases where task constraints completely overwhelm all system-specific constraints, might we expect the natural emergence of alignment at the single-unit level.

Returning to grid cells, since they do not generically arise in networks trained to path integrate, path integration is not a sufficiently constraining task. Theoretical work on grid cell representations [22, 61, 45] suggests additional critical features of the code: an exponentially large coding range and robustness/intrinsic error correction, both of which translate into the problem of packing and maximally separating a large set of coding states into a compact space [61]. We hypothesize that the following key properties of the grid cell code may form a biologically relevant sufficient set for their emergence: 1) non-negative activity; 2) path integrating (PI) code that is translation invariant [23, 9, 12]; 3) exponential representational capacity [8, 61, 44]; 4) intrinsic error-correcting capabilities [8, 61]; and 5) uniformly distributed (whitened) information across cells. Several of these are general properties of neural codes, and could increase the ability of ANN models to make de novo rather than post hoc predictions. For recent promising work, see [17].

First-principles continuous attractor network (CAN) models of grid cells made several novel predictions subsequently confirmed in experiments [39]: the invariance of cell-cell relationships acrossenvironments and behavioral states (constituting, in the terminology of machine learning, “far out of distribution” predictions) that define an invariant toroidal attractor manifold [72, 64, 26, 25], grid-like patterning in the cortical sheet [29], and many others that remain to be tested. Deep learning-based models should be held to similar standards.

In sum, ANN models of the brain that reproduce specific tuning curves should not center their claimed achievement on producing the curves if these are used as implicit or explicit parts of the training target (given the expressive power of deep networks, it is no revelation that training them to generate a given tuning will in fact succeed), but rather should characterize the conditions under which the particular tuning does and does not emerge, to consider which inductive biases are critical, and to explicitly state what principles and de novo predictions can be extracted from the models.

## 9 Acknowledgements

This work was supported by the ONR, the NSF, the Simons Foundation through the SSCGB program, and the HHMI through the Faculty Scholars Program. MK was supported by the MathWorks Science Fellowship.

## 10 Checklist

1. For all authors…
  a. Do the main claims made in the abstract and introduction accurately reflect the paper’s contributions and scope?
  b. Did you describe the limitations of your work?
  c. Did you discuss any potential negative societal impacts of your work? We do not feel our paper has potential negative societal impacts.
  d. Have you read the ethics review guidelines and ensured that your paper conforms to them?
2. If you are including theoretical results…
  a. Did you state the full set of assumptions of all theoretical results?
  b. Did you include complete proofs of all theoretical results?
3. If you ran experiments…
  a. Did you include the code, data, and instructions needed to reproduce the main experimental results (either in the supplemental material or as a URL)?
  b. Did you specify all the training details (e.g., data splits, hyperparameters, how they were chosen)?
  c. Did you report error bars (e.g., with respect to the random seed after running experiments multiple times)?
  d. Did you include the total amount of compute and the type of resources used (e.g., type of GPUs, internal cluster, or cloud provider)? We haven’t yet had time to collect this information, but we will add this information to the final version if accepted.
4. If you are using existing assets (e.g., code, data, models) or curating/releasing new assets…
  a. If your work uses existing assets, did you cite the creators?
  b. Did you mention the license of the assets?
  c. Did you include any new assets either in the supplemental material or as a URL?
  d. Did you discuss whether and how consent was obtained from people whose data you’re using/curating?
  e. Did you discuss whether the data you are using/curating contains personally identifiable information or offensive content?
5. If you used crowdsourcing or conducted research with human subjects…
  a. Did you include the full text of instructions given to participants and screenshots, if applicable?
  b. Did you describe any potential participant risks, with links to Institutional Review Board (IRB) approvals, if applicable?
  c. Did you include the estimated hourly wage paid to participants and the total amount spent on participant compensation?

## A Position Encodings

Suppose we sample a sequence of positions *x*_0_, …, *x*_*T*_ ∈ ℝ^2^ and a sequence of velocities *v*_1_, …, *v*_*T*_ ∈ ℝ^2^, where *x*_*t*_ = *x*_*t*−1_ + *v*_*t*_. We want to train the networks in a supervised manner to predict (a possible encoding of) their position. We used the below encodings as different regression targets. For some encodings that required place cell populations, we denote the number of place cells *N*_*p*_ and denote their locations 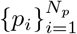, sampled uniformly at random within the 2.2 m × 2.2 m arena.

- **“Low-dimensional” Cartesian:** *x*_*t*_
- **“High-dimensional” Cartesian:** Let 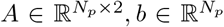 have entries sampled i.i.d. from *Uniform*(−1, 1). The target is a vector in 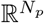 given by:

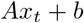

- **Polar:** Let (*r*_*t*_, *θ*_*t*_) denote the polar form of the agent’s position *x*_*t*_. The target is a vector in 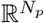, half comprised of “distance” units and half comprised of “direction” units. Let 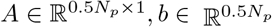 have entries sampled i.i.d. from *Uniform*(−1, 1); the distance cells have activites: 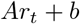 Let 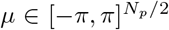 have entries sampled i.i.d. uniformly at random. The direction cells have entries given by von-Mises-like bumps:

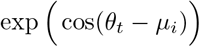
- **Gaussian:** A vector in 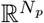 whose entries are given by:

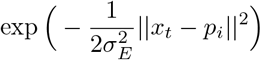

- **equi-norm Difference of Gaussians:** A vector in 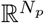 whose entries are given by:

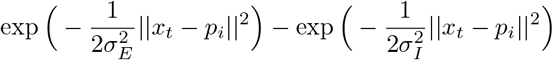

- **Difference of Softmaxes:** A vector in 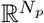 whose entries are given by:

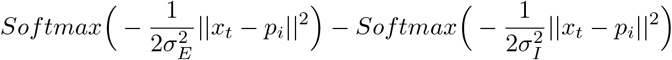

- **Softmax of Differences:** A vector in 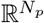 whose entries are given by:

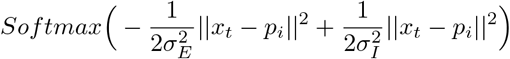

## B Grid Scores and Grid Cell Thresholds

What qualifies as a grid cell? The most commonly used method of quantifying grid cells is via the “grid score”, which functions by binning neural activity into rate maps using spatial position, applying an adaptive smoother, then taking a circular sample of the autocorrelation centered on the central peak and comparing it to rotated versions of the same circular sample. The 60 ° grid score is specifically given by:

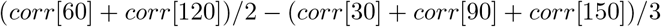

We used the same grid scorer implementation used by [3] (<https://github.com/deepmind/grid-cells/blob/master/scores.py>), [59] (https://github.com/ganguli-lab/grid-pattern-formation/blob/master/scores.py) and [47].

What score is sufficient to qualify as a grid score? Experimentalists have used thresholds of 0.3 [58] and 0.349 [11] on biological neurons, whereas computationalists have used 0.3 [47] and 0.37 [3, 59] on artificial neurons. We found that for artificial neurons, these thresholds are far too low (Fig. 8); ANN units with grid scores > 0.4, and even as high as 1.3, often look nothing like grid cells. This is because the grid score looks for 60° rotational symmetry, and while grid cells are indeed symmetric, so are many other rate maps.

**Figure 8:**
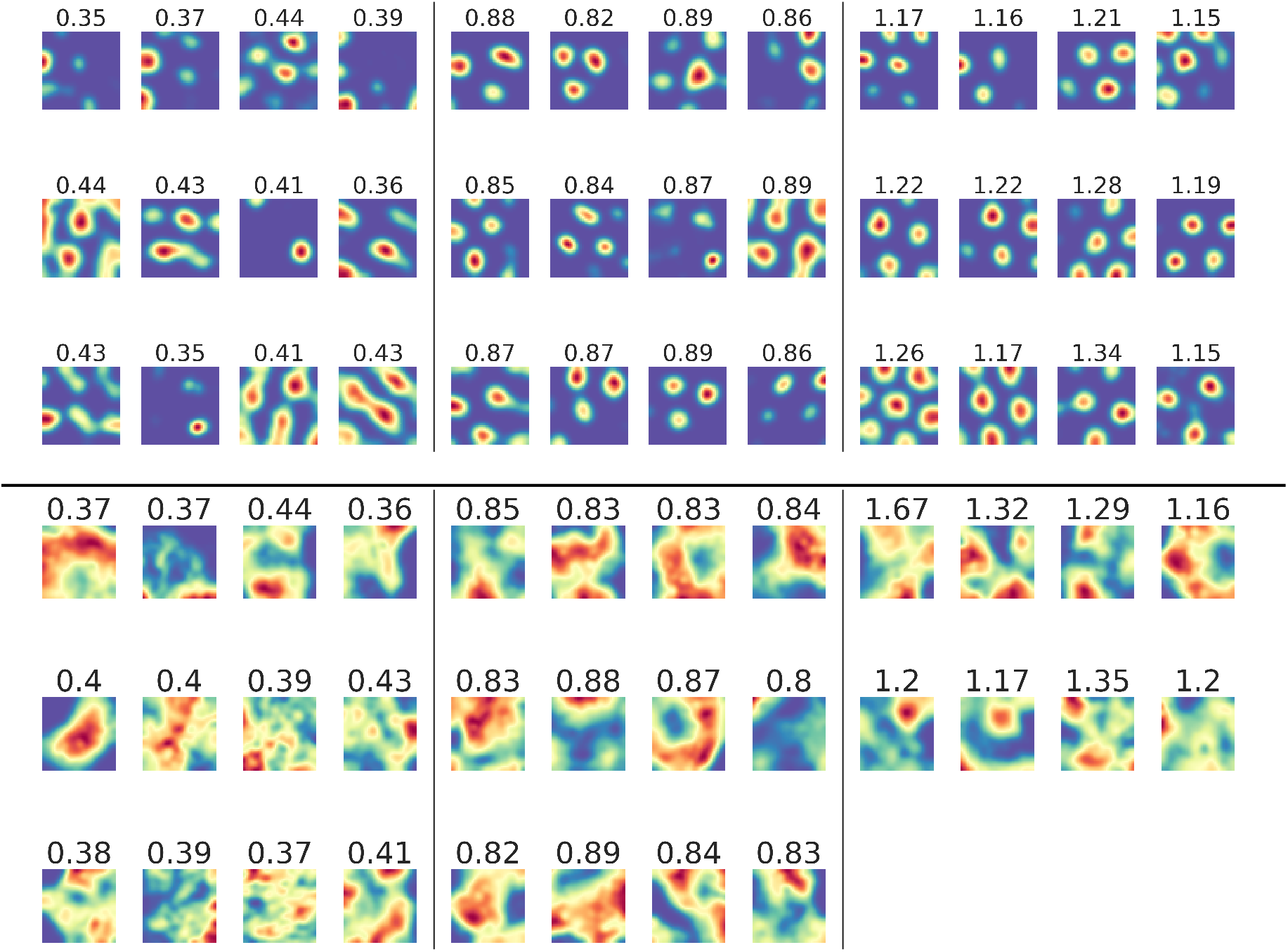
Grid scoring is the dominant method to identify grid cells, but we found it performs inadequately at identifying grid cells. Top: example rate maps from low position decoding error Difference-of-Softmaxes (DoS) networks three grid score ranges: [0.35, 0.45), [0.8, 0.9), [1.15, ∞). Bottom: example rate maps from low position decoding error Difference-of-Gaussians (DoG) networks. We considered three grid score thresholds: 0.3 (used by some experimentalists), 0.8 (low probability of finding grid cells), 1.15 (decent probability of finding grid cells).

## C Number of Bins for Computing Rate Maps

The first step in computing grid scores is determining the number of bins to use to compute rate maps. The original experimental work used 5 cm x 5 cm bins [30]. Since the square arena used in these experiments is 2.2 m by 2.2 m (same as [3, 59, 47]), the number of bins should be 44 x 44. Due to inconsistencies in the number of bins previously used, we checked what effect, if any, the number of bins has on the distribution of grid scores. We found that 20×20, 32×32 and 44×44 appears to have little to no differences (Fig. 9), so we chose to use 44 x 44 bins to be consistent with experimentalists.

**Figure 9:**
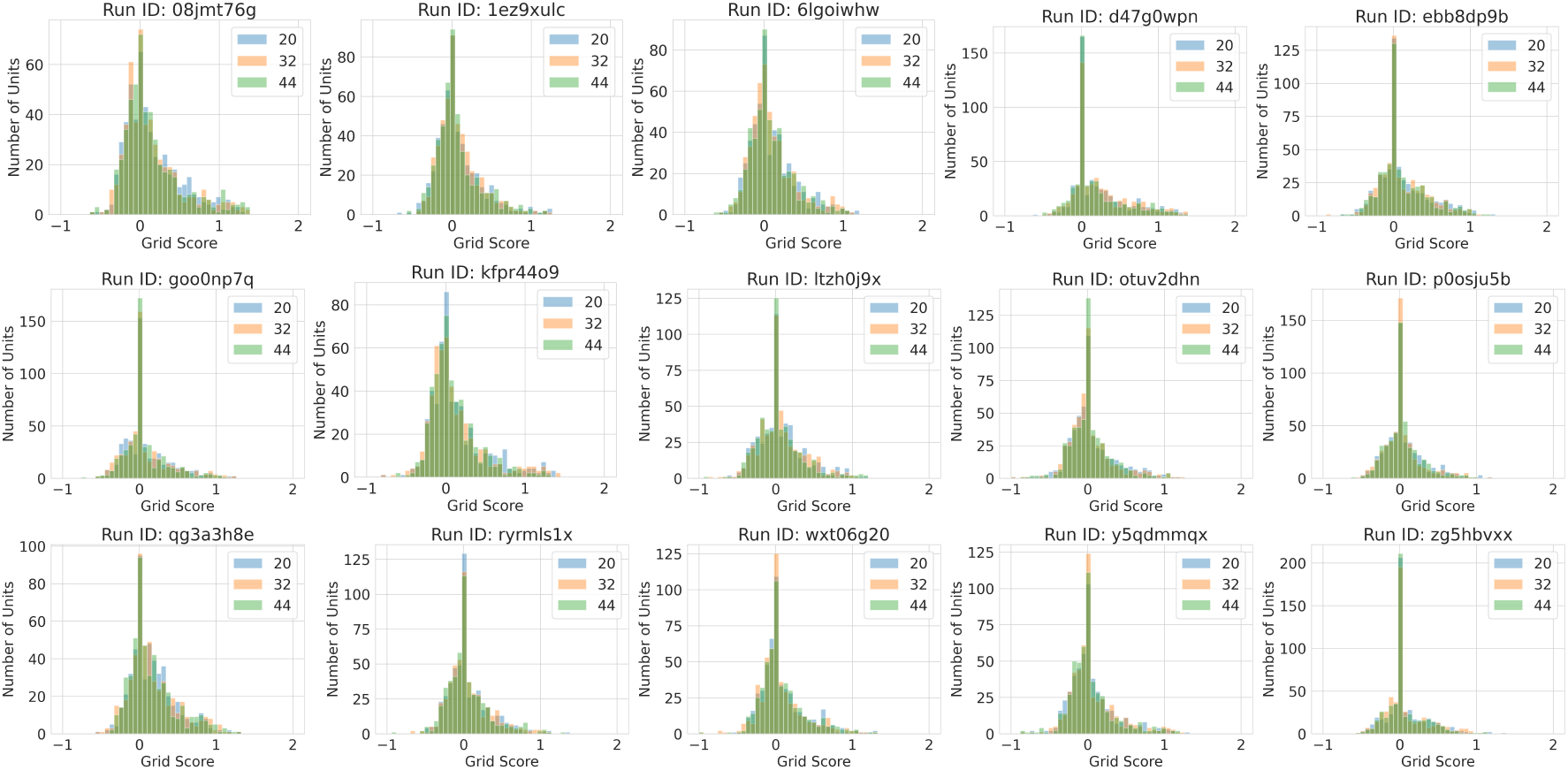
Grid score distributions do not differ as a function of number of bins: 400 (20 x 20; blue), 1024 (32 x 32; orange), 1936 (44 x 44; green).

## D Place cell spatial autocorrelation

In this section, we will derive the form of the place cell correlation function. When the spatial tuning curve is a difference-of-Gaussians, the correlation function is also a Difference-of-Gaussians, albeit with different parameters. This calculation is performed in d dimensions. In simulations, *d* = 2. Consider the spatial tuning curve:

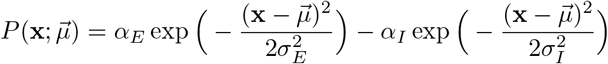

So the correlation function of **x**_**1**_ and **x**_**2**_ with **∆x** = **x**_**1**_ − **x**_**2**_ (assuming that place cell centers are distributed isotropically and can be integrated over) is given by:

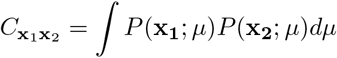

Simplifying the above expression, using the identity: 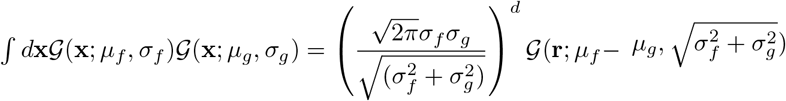, where 𝒢 (x; *μ*, σ) = exp (−(x − *μ*)^2^/2σ^2^)

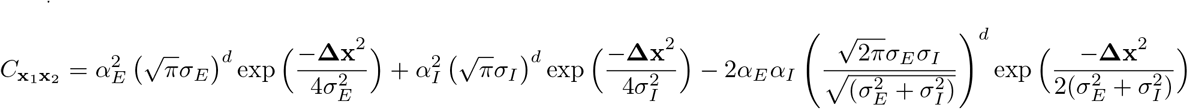

The Fourier transform of this quantity, 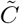 can be calculated using the identity

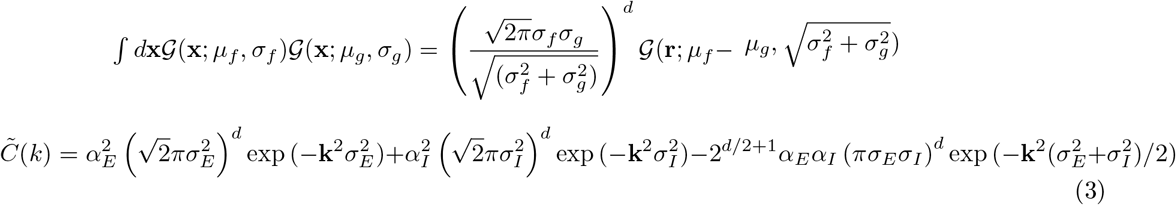

Using this equation, we numerically solve for the maximum *k*^*^.

## E DoG and Narrow DoG Sweeps

We evaluated non-equinorm Difference-of-Gaussian readouts, with activations given by:

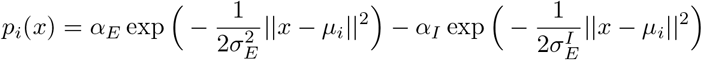

The analytics tell us that the ratio *α*_*E*_/*α*_*I*_ should determine whether the Fourier annulus radius > 1 and thus whether grid-like representations might emerge. Over a broad sweep of *α*_*E*_/*α*_*I*_ ratio values, we found that all networks’ grid score distributions were dominated by filtered-then-ReLU-thresholded noise ∼*Uniform*(1, −1), which were in turn dominated by ideal-width Difference-of-Softmaxes readouts (Fig. 10). This shows that most DoG readouts will fail to produce lattices.

**Figure 10:**
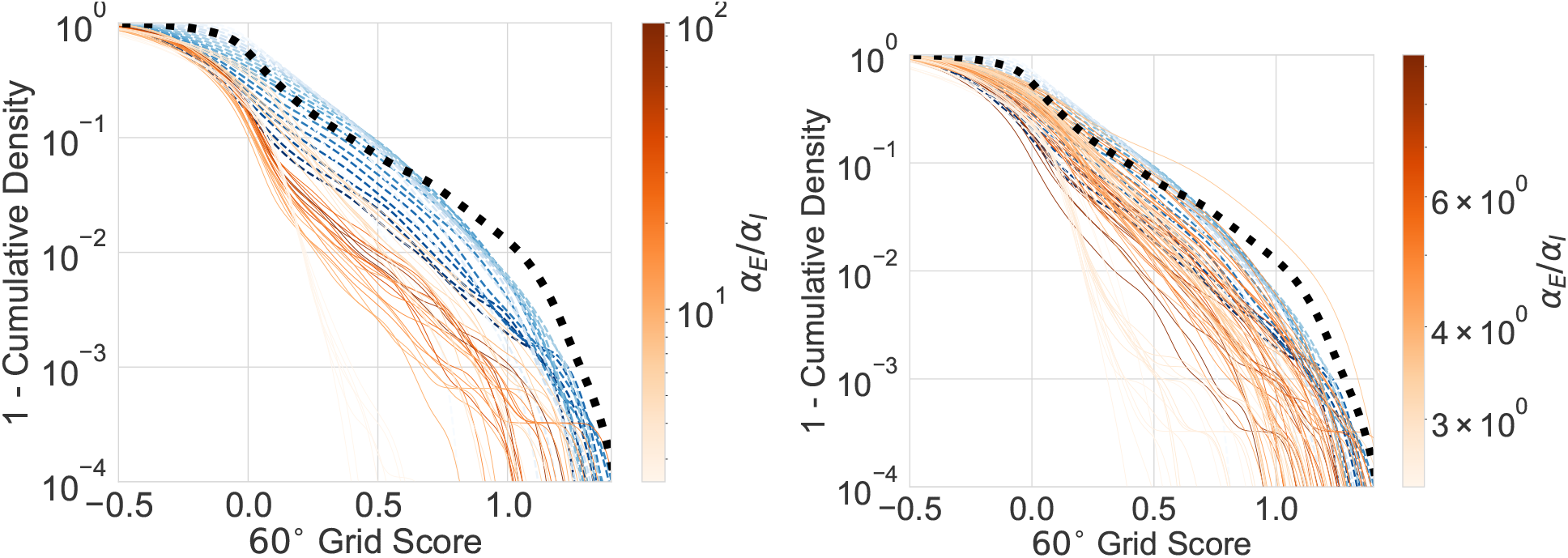
Left: Fig. 4 from main text. A broad sweep of DoG hyperparameters show that DoG (red) obtains lower grid score distributions than filtered-and-thresholded noise (blue) and than Difference-of-Softmaxes (black). Right: A narrow sweep of DoG hyperparameters in the “feasible region” of Fig. 4c show that almost all DoG (red) obtains lower or comparable grid score distributions than filtered-and-thresholded noise (blue), and lower than Difference-of-Softmaxes (black). Note: Colorbars are different between left and right.

Next, we ran a much narrower sweep of *α*_*E*_/*α*_*I*_ ratio values in the range (2.5, 10), which [59] predicts should robustly yield lattices. We found that most networks’ grid score distributions were dominated by or indistinguishable from filtered-then-ReLU-thresholded noise ∼*Uniform*(1, −1), which were in turn dominated by ideal-width Difference-of-Softmaxes readouts (Fig. 10). One ratio *α*_*E*_/*α*_*I*_ = 3.5417 appeared to outperform DoS, but this was due to a single lucky seed (Fig. 4e). This shows that, even when theory predicts the formation of lattices, most DoG readouts will fail to produce lattices.

## F Sweeps

### F.1 Cartesian (Low Dimensional)

~~~
method : grid
metric :
  goal : minimize
  name : pos_decoding_err
parameters :
  Ng :
      values:
        − 1024
  Np :
      values :
       − 2
activation :
       values :
         − relu
         − tanh
         − sigmoid
         − linear
batch_size:
       values:
         − 200
bin_side_in_m :
       values:
         − 0.05
box_height_in_m:
       values:
         − 2.2
box_width_in_m:
       values:
         − 2.2
initializer :
       values:
         − glorot_uniform
         − glorot_normal
         − orthogonal
is_periodic :
       values :
         − false
learning_rate:
       values :
         − 0.0001
n_epochs :
       values :
         − 20
n_grad_steps_per_epoch :
       values :
         − 10000
n_place_fields_per_cell :
       values :
         − 1
optimizer :
       values :
         − sgd
         − adam
         − rmsprop
place_cell_rf:
       values :
         − 0
place_field_loss:
       values :
         − mse
place_field_normalization :
       values :
         − none
place_field_values :
       values :
         − cartesian
readout_dropout :
       values :
         − 0
         − 0.5
recurrent_dropout :
       values :
         − 0
         − 0.5
rnn_type :
       values :
         − RNN
         − LSTM
         − UGRNN
         − GRU
seed :
       values :
         − 0
         − 1
         − 2
sequence_length :
       values :
         − 20
sur round_scale :
       values :
         − 1
weight_decay :
       values :
         − 0
         − 0.0001
~~~

### F.2 Cartesian (Low Dimensional, Random)

~~~
method : random
metric :
  goal : minimize
   name: pos_decoding_err
parameters :
activation :
       values : [
         ‘ relu ‘,
         ‘ tanh ‘,
         ‘ sigmoid ‘,
         ‘ linear ‘,
]
batch_size :
       values : [30, 60, 90, 120, 150, 180]
bin_side_in_m :
       values : [0.05 ]
box_height_in_m:
       values : [2.2 ]
box_width_in_m:
       values : [2.2 ]
initializer:
       values : [
         ‘ glorot_uniform ‘,
         ‘ glorot_normal ‘,
         ‘ orthogonal ‘,
]
is_periodic :
       values : [False]
learning_rate :
       values : [0.005, 0. 001, 0. 0005, 0. 0001]
n_epochs :
       values : [1, 4, 8, 12, 16 ]
n_grad_steps_per_epoch :
       values : [10000]
n_place_fields_per_cell :
       values : [1]
Ng:
       values : [102 4 ]
Np:
       values : [2 ]
optimizer :
       values : [
         ‘adam’,
         ‘ rmsprop ‘,
]
place_field_loss:
       values  : [
         ‘mse ‘,
]
place_field_values :
      values  : [
         ‘ cartesian ‘,
]
place_field_normalization :
      values  : [
         ‘ none ‘,
]
place_cell_rf:
      values  : [
0.
]
readout_dropout :
      values  : [0., 0. 05, 0. 1, 0. 2, 0. 5 ]
recurrent_dropout :
      values  : [0., 0. 05, 0. 1, 0. 2, 0. 5 ]
rnn_type :
      values  : [
         ‘RNN’,
         ‘LSTM’,
         ‘UGRNN’,
         ‘GRU’,
]
seed :
      values  : [0, 1, 2 ]
sequence_length :
      values  : [2 0, 25, 30, 35, 4 0]
sur round_scale :
      values  : [1. ]
weight_decay :
      values  : [0., 0. 0001, 0. 0005,0.001, 0. 005, 0. 01]
~~~

### F.3 Cartesian (High Dimensional)

~~~
method : grid
metric :
goal : minimize
name : pos_decoding_err
parameter s :
Ng:
      values  :
         − 1024
Np:
      values :
         − 512
activation :
      values  :
         − relu
         − tanh
         − sigmoid
         − linear
batch_size:
      values  :
         − 200
bin_side_in_m :
      values  :
         − 0.05
box_height_in_m:
      values  :
         − 2.2
box_width_in_m:
      values  :
         − 2.2
i n i t i a l i z e r :
      values  :
         − glorot_uniform
         − glorot_normal
         − orthogonal
i s_periodic:
      values  :
         − false
learning_rate:
      values  :
         − 0.0001
n_epochs :
      values  :
         − 20
n_grad_steps_per_epoch :
      values  :
         − 1000
n_plac e_fields_pe r_cell :
      values  :
         − 1
optimizer:
      values  :
         − adam
place_cell_rf:
      values  :
         − 0
p l a c e_field_loss:
      values  :
         − mse
place_field_normalization :
      values  :
         − none
place_field_      values  :
      values  :
         − high_dim_cartesian
readout_dropout :
      values  :
         − 0
         − 0. 5
recurrent_dropout :
      values  :
         − 0
         − 0. 5
rnn_type :
      values  :
         − RNN
         − LSTM
         − UGRNN
         − GRU
seed :
      values  :
         − 0
         − 1
         − 2
sequence_length :
      values  :
         − 20
sur round_scale :
      values  :
         − 0
weight_decay :
      values  :
         − 0
         − 0.0001
~~~

### F.4 Cartesian (High Dimensional, Random)

~~~
method : random
metric :
  goal : minimize
  name : pos_decoding_err
parameters :
activation :
      values: [
         ‘ relu ‘,
         ‘ tanh ‘,
         ‘ sigmoid ‘,
         ‘ linear ‘,
]
batch_size :
      values: [3 0, 60, 90, 120, 150, 180]
bin_side_in_m :
      values: [0. 05 ]
box_height_in_m :
      values: [2 . 2 ]
box_width_in_m :
      values: [2 . 2 ]
initializer :
      values: [
         ‘ glorot_uniform ‘,
         ‘ glorot_normal ‘,
         ‘ orthogonal ‘,
]
is _periodic :
      values : [False ]
learning _rate :
      values : [0. 005, 0. 001, 0. 0005, 0. 0001]
n_epochs :
      values : [1, 4, 8, 12, 16 ]
n_grad_steps_per_epoch :
      val u e s : [1000]
n_place_fields_per_cell :
      values: [1]
  Ng :
      values: [102 4 ]
  Np :
      values: [12 8, 256, 512 ]
optimizer :
      values: [
        ‘ adam ‘,
        ‘ rmsprop ‘,
]
place _field _loss:
      values: [
        ‘ mse ‘,
]
place _field _values:
      values: [
        ‘ high_dim_cartesian ‘
]
place _field _normalization :
      values: [
        ‘ none ‘,
]
place _cell _r f :
      values: [
    0. 0,
]
recurrent_dropout :
      values: [0., 0. 05, 0. 1, 0. 2, 0. 5 ]
readout_dropout :
      values: [0., 0. 05, 0. 1, 0. 2, 0. 5 ]
rnn_type :
      values: [
        ‘RNN’,
        ‘LSTM’,
        ‘UGRNN’,
        ‘GRU’,
]
seed :
      values: [0, 1, 2 ]
sequence_length :
      values: [2 0, 25, 30, 35, 4 0]
surround_scale :
      values: [
  0.
]
weight_decay :
      values: [0., 0. 0001, 0. 0005, 0. 001, 0. 005, 0. 01]
~~~

### F.5 Polar (High Dimensional)

~~~
method : g r i d
metric :
  goal : minimize
  name : pos_decoding_err parameters :
Ng :
      values:
        − 1024
Np :
      values:
        − 512
activation :
      values:
        − relu
        − tanh
        − linear
        − sigmoid
batch_size :
      values:
        − 200
bin_side_in_m :
      values:
        − 0. 05
box_height_in_m :
      values:
        − 2 . 2
box_width_in_m : values:
        − 2 . 2
initializer :
      values:
        − glorot_uniform
        − glorot_normal
        − orthogonal i s _periodic :
      values:
        − false
learning _r a te :
      values:
        − 0. 0001
n_epochs :
      values:
        − 20
n_grad_steps_per_epoch :
      values:
        − 1000
n_place_fields_per_cell :
      values:
        − 1
optimizer :
      values:
        − adam
place _cell _r f :
      values:
        − 0
place _field _loss:
      values:
        − mse
place _field _normalization :
      values:
        − none
place _field _values:
      values:
        − high_dim_polar readout_dropout :
      values:
        − 0
        − 0. 5
recurrent_dropout :
      values:
        − 0
        − 0. 5
rnn_type :
      values:
        − RNN
        − LSTM
        − UGRNN
        − GRU
seed :
      values:
        − 0
        − 1
        − 2
sequence_length :
      values:
        − 20
surround_scale :
      values:
        − 0
weight_decay :
      values:
        − 0
        − 0. 0001
~~~

### F.6 Polar (High Dimensional, Random)

~~~
method : random
metric :
  goal : minimize
  name : pos_decoding_err
parameters :
activation :
     values: [
        ‘ relu ‘,
        ‘ tanh ‘,
        ‘ linear ‘,
        ‘ sigmoid ‘,
]
batch_size :
    values: [3 0, 60, 90, 120, 150, 180]
bin_side_in_m :
    values: [0. 05 ]
box_height_in_m :
    values: [2 . 2 ] box_width_in_m :
    values: [2 . 2 ]
initializer :
    values: [
      ‘ glorot_uniform ‘,
      ‘ glorot_normal ‘,
      ‘ orthogonal ‘,
]
i s _periodic :
    values:
     [False ]
learning _r a te :
    values: [0. 005, 0. 001, 0. 0005, 0. 0001]
n_epochs :
    values: [1, 4, 8, 12, 16 ]
n_grad_steps_per_epoch :
    values: [5 000]
n_place_fields_per_cell :
    values: [1]
 Ng :
    values: [102 4 ]
 Np :
    values: [12 8, 256, 512 ]
optimizer :
    values: [
      ‘ adam ‘,
      ‘ rmsprop ‘,
]
place _field _loss:
    values: [
      ‘ mse ‘,
]
place _field _values:
    values: [
      ‘ high_dim_polar ‘,
]
place _field _normalization :
    values: [
      ‘ none ‘,
]
place _cell _r f :
    values: [
    0.
  ]
readout_dropout :
    values: [0., 0. 05, 0. 1, 0. 2, 0. 5 ]
recurrent_dropout :
    values: [0., 0. 05, 0. 1, 0. 2, 0. 5 ]
rnn_type :
    values: [
      ‘RNN’,
      ‘LSTM’,
      ‘UGRNN’,
      ‘GRU’,
  ]
seed :
    values: [0, 1, 2 ]
sequence_length :
    values: [2 0, 25, 30, 35, 4 0]
surround_scale :
    values: [0. ]
weight_decay :
    values: [0., 0. 0001, 0. 0005, 0. 001, 0. 005, 0. 01]
~~~

### F.7 Gaussian Place Cells

~~~
method : g r i d
metric :
  goal : minimize
  name : pos_decoding_err
parameters :
  Ng :
    values:
      − 1024
  Np :
    values:
      − 512
activation :
    values:
      − linear
      − relu
      − tanh
      − sigmoid
batch_size :
    values:
      − 200
bin_side_in_m :
    values:
      − 0. 05
box_height_in_m :
    values:
      − 2 . 2
box_width_in_m :
    values:
     − 2 . 2
initializer :
    values:
      − glorot_uniform
i s _periodic :
    values:
      − f a l s e
learning _r a te :
    values:
      − 0. 0001
n_epochs :
    values:
     − 20
n_grad_steps_per_epoch :
    values:
     − 10000
n_place_fields_per_cell :
    values:
      − 1
optimizer :
    values:
      − adam
place _cell _r f :
    values:
      − 0. 08
      − 0. 1
      − 0. 12
      − 0. 14
      − 0. 16
      − 0. 2
      − 0. 24
      − 0. 28
place _field _loss:
    values:
      − crossentropy
place _field _normalization :
    values:
      − global
place _field _values:
    values:
      − gau s s i an readout_dropout :
    values:
      − 0
      − 0. 5
recurrent_dropout :
    values:
      − 0
      − 0. 5
rnn_type :
    values:
      − RNN
      − LSTM
      − UGRNN
      − GRU
seed :
    values:
      − 0
      − 1
sequence_length :
    values:
      − 20
surround_scale :
    values:
      − 1
weight_decay :
    values:
      − 0. 0001
~~~

### F.8 Gaussian Place Cells (Random)

~~~
method : random
metric :
  goal : minimize
  name : pos_decoding_err
parameters :
activation :
     values: [
        ‘ linear ‘,
        ‘ relu ‘,
        ‘ tanh ‘,
        ‘ sigmoid ‘,
]
batch_size :
    values: [3 0, 60, 90, 120, 150, 180]
bin_side_in_m :
    values: [0. 05 ] box_height_in_m :
    values: [2 . 2 ] box_width_in_m : values: [2 . 2 ]
initializer :
    values: [
      ‘ glorot_uniform ‘,
      ‘ glorot_normal ‘,
      ‘ orthogonal ‘,
]
i s _periodic :
    values: [False ] learning _r a te :
    values: [0. 005, 0. 001, 0. 0005, 0. 0001]
n_epochs :
    values: [1, 4, 8, 12, 16 ]
n_grad_steps_per_epoch :
    values: [10000]
n_place_fields_per_cell :
    values: [1. 0,
]
Ng :
    values: [102 4 ]
Np :
    values: [12 8, 256, 512 ] optimizer :
    values: [‘ adam ‘,
      ‘ rmsprop ‘
]
place _field _loss:
    values: [
      ‘ crossentropy ‘,
]
place _field _values:
    values: [
      ‘ gaussian ‘,
]
place _field _normalization :
    values: [
      ‘ g l obal ‘,
]
place _cell _r f :
    values: [
      0. 08,
      0. 10,
      0. 12,
      0. 14,
      0. 16,
      0. 2 0,
      0. 2 4,
      0. 2 8,
      0. 3 2,
      0. 3 6,
      0. 4 0,
]
readout_dropout :
    values: [0., 0. 05, 0. 1, 0. 2, 0. 5 ]
recurrent_dropout :
   values: [0., 0. 05, 0. 1, 0. 2, 0. 5 ]
rnn_type :
   values: [
     ‘RNN’,
     ‘LSTM’,
     ‘UGRNN’,
     ‘GRU’,
]
seed :
   values: [0, 1, 2 ]
sequence_length :
   values: [2 0, 25, 30, 35, 4 0]
surround_scale :
   values: [1. ]
weight_decay :
   values: [0., 0. 0001, 0. 0005, 0. 001, 0. 005, 0. 01]
~~~

### F.9 Difference-of-Gaussians Place Cells

~~~
method : g r i d
metric :
  goal : minimize
  name : pos_decoding_err
parameters :
  Ng :
   values:
     − 1024
  Np :
   values:
     − 512
activation :
   values:
     − relu batch_size :
   values:
    − 200
bin_side_in_m :
   values:
     − 0. 05
box_height_in_m :
   values:
     − 2 . 2
box_width_in_m :
   values:
     − 2 . 2
initializer :
   values:
     − glorot_uniform i s _periodic :
   values:
     − f a l s e
learning _r a te :
   values:
     − 0. 0001
n_epochs :
  values:
     − 20
n_grad_steps_per_epoch :
   values:
     − 10000
n_place_fields_per_cell :
   values:
     − 1
optimizer :
   values:
     − adam
place _cell _r f :
   values:
     − 0. 05
     − 0. 1
     − 0. 15
     − 0. 2
     − 0. 3
     − 0. 4
     − 0. 5
place _field _loss:
  values:
    − crossentropy
place _field _normalization :
     values:
       − global
place _field _values:
     values:
       − tr u e _d i f f e re n c e _o f _ga u s s i a n s
readout_dropout :
     values:
       − 0
       − 0. 5
recurrent_dropout :
     values:
       − 0
       − 0. 5
rnn_type :
     values:
       − RNN
       − LSTM
       − UGRNN
       − GRU
seed :
     values:
       − 0
       − 1
       − 2
sequence_length :
     values:
       − 20
surround_scale :
     values:
       − 1. 5
       − 2
       − 2 . 5
       − 3
       − 4
       − 5
       − 6
weight_decay :
     values:
       − 0. 0001
~~~

### F.10 Difference-of-Gaussians Place Cells (Random)

~~~
method : random
metric :
  goal : minimize
  name : pos_decoding_err
parameters :
  activation :
    values: [
      ‘ linear ‘,
      ‘ relu ‘,
      ‘ tanh ‘,
      ‘ sigmoid ‘,
]
batch_size :
    values: [3 0, 60, 90, 120, 150, 180]
bin_side_in_m :
    values: [0. 05 ]
box_height_in_m :
    values: [2 . 2 ]
box_width_in_m :
    values: [2 . 2 ]
initializer :
    values: [
      ‘ glorot_uniform ‘,
      ‘ glorot_normal ‘,
      ‘ orthogonal ‘,
]
i s _periodic :
    values: [False ]
learning _r a te :
    values: [0. 005, 0. 001, 0. 0005, 0. 0001]
n_epochs :
    values: [1, 4, 8, 12, 16 ]
n_grad_steps_per_epoch :
    values: [10000]
n_place_fields_per_cell : values: [1]
Ng :
    values: [102 4 ]
Np :
    values: [12 8, 256, 512 ]
optimizer :
    values: [
      ‘ adam ‘,
      ‘ rmsprop ‘
]
place _field _loss:
    values: [
     ‘ crossentropy ‘,
]
place _field _values:
    values: [
      ‘ true_difference_of_gaussians ‘
]
place _field _normalization :
    values: [
      ‘ g l obal ‘,
]
place _cell _r f :
    values: [
     0. 08,
     0. 10,
     0. 12,
     0. 14,
     0. 16,
     0. 2 0,
     0. 2 4,
     0. 2 8,
     0. 3 2,
     0. 3 6,
     0. 4 0,
]
recurrent_dropout :
    values: [0., 0. 05, 0. 1, 0. 2, 0. 5 ]
readout_dropout :
    values: [0., 0. 05, 0. 1, 0. 2, 0. 5 ]
rnn_type :
    values: [‘RNN’,
      ‘LSTM’,
      ‘UGRNN’,
      ‘GRU’,
]
seed :
    values: [0, 1, 2 ]
sequence_length :
    values: [2 0, 25, 30, 35, 4 0]
surround_scale :
    values: [
      1. 5,
      2 .,
      2 . 5,
      3 .,
      3 . 5,
      4 .,
      4 . 5,
      5 .,
      5 . 5,
      6 .,
]
weight_decay :
    values: [0., 0. 0001, 0. 0005, 0. 001, 0. 005, 0. 01]
~~~

### F.11 Difference-of-Softmaxes Place Cells

~~~
method : g r i d
metric :
  goal : minimize
  name : pos_decoding_err
parameters :
 Ng :
   values:
     − 1024
Np :
   values:
     − 512
activation :
   values:
     − relu
     − tanh batch_size :
   values:
     − 200
bin_side_in_m :
   values:
     − 0. 05
box_height_in_m :
   values:
     − 2 . 2
box_width_in_m :
   values:
     − 2 . 2
initializer :
   values:
     − glorot_uniform i s _periodic :
   values:
     − f a l s e
learning _r a te :
   values:
     − 0. 0001
n_epochs :
   values:
     − 20
n_grad_steps_per_epoch :
   values:
     − 10000
n_place_fields_per_cell :
   values:
     − 1
optimizer :
  values:
   − adam
place _cell _r f :
  method : random
    − 0. 08
    − 0. 1
    − 0. 12
    − 0. 14
    − 0. 16
    − 0. 2
    − 0. 24
    − 0. 28
place _field _loss:
  values:
    − crossentropy
place _field _normalization :
  values:
    − global
place _field _values:
  values:
    − gau s s i an readout_dropout :
   values:
    − 0
    − 0. 5
recurrent_dropout :
   values:
   − 0
   − 0. 5
rnn_type :
   values:
     − RNN
     − LSTM
   − UGRNN
   − GRU
seed :
  values:
   − 0
   − 1
sequence_length :
  values:
   − 20
surround_scale :
  values:
   − 1
weight_decay :
  values:
   − 0. 0001
~~~

### F.12 Difference-of-Softmaxes Place Cells (Random)

~~~
method : random
me t r i c :
  go al : minimize
  name : pos_decoding_err
parameter s :
  a c t i v a t i o n :
   va lue s : [
    ‘ linear ‘,
    ‘ r e lu ‘,
    ‘ tanh ‘,
    ‘ sigmoid ‘,
]
batch_s ize :
   va lue s : [3 0, 60, 90, 120, 150, 180]
bin_side_in_m :
   va lue s : [0. 05 ]
box_height_in_m:
   va lue s : [2 . 2 ]
box_width_in_m:
   va lue s : [2 . 2 ]
i n i t i a l i z e r :
   va lue s : [
    ‘ glorot_uniform ‘,
    ‘ glorot_normal ‘,
    ‘ or thogonal ‘,
]
i s_periodic :
   va lue s : [Fal s e ]
l e a rning_r a t e :
   va lue s : [0. 005, 0. 001, 0. 0005, 0. 0001]
n_epochs :
   va lue s : [2 0]
n_grad_steps_per_epoch :
   va lue s : [10000]
n_plac e_f i e lds_pe r_c e l l :
   va lue s : [
1. 0,
]
Ng:
   value s : [102 4 ]
Np:
   value s : [12 8, 256, 512 ]
opt imi z e r :
   value s : [
    ‘adam’,
    ‘ rmsprop ‘,
]
place_f i e l d_l o s s :
   va lue s : [
     ‘ c ros s ent ropy ‘,
]
place_f i e ld_va lue s :
   va lue s : [
     ‘ di f f e r enc e_o f_g aus s i ans ‘,
]
place_f i e ld_no rma l i z a t i on :
   va lue s : [
     ‘ global ‘,
]
place_c e l l_r f :
   va lue s : [
    0. 08,
    0. 10,
    0. 12,
    0. 14,
    0. 16,
    0. 2 0,
    0. 2 4,
    0. 2 8,
    0. 3 2,
    0. 3 6,
    0. 4 0,
    0. 4 4,
    0. 4 8,
]
readout_dropout :
   va lue s : [0., 0. 05, 0. 1, 0. 2, 0. 5 ]
recur rent_dropout :
46
   va lue s : [0., 0. 05, 0. 1, 0. 2, 0. 5 ]
rnn_type :
   va lue s : [
    ‘RNN’,
    ‘LSTM’,
    ‘UGRNN’,
    ‘GRU’,
]
seed:
   va lue s : [0, 1, 2 ]
sequence_length :
   va lue s : [2 0, 25, 30, 35, 4 0]
sur round_scale :
   va lue s : [
    1. 5,
    2 .,
    2 . 5,
    3 . 0,
    3 . 5,
    4 . 0,
    4 . 5,
    5 . 0,
    5 . 5,
    6 . 0,
]
weight_decay :
   va lue s : [0., 0. 0001, 0. 0005, 0. 001, 0. 005, 0. 01]
~~~

### F.13 Softmax-of-Differences Place Cells

~~~
method : g r id
me t r i c :
  go al : minimize
  name : pos_decoding_err
parameter s :
Ng:
   v a lue s :
     − 1024
Np:
   v a lue s :
     − 512
a c t i v a t i o n :
   va lue s :
     − relu
batch_s ize :
   va lue s :
     − 200
bin_side_in_m :
   va lue s :
    − 0. 05
box_height_in_m:
47
   va lue s :
    − 2 . 2
box_width_in_m:
   va lue s :
    − 2 . 2
i n i t i a l i z e r :
va lue s :
    − glorot_uniform
i s_periodic :
   va lue s :
    − f a l s e
l e a rning_r a t e :
   va lue s :
    − 0.0001
n_epochs :
   va lue s :
    − 20
n_grad_steps_per_epoch :
   va lue s :
    − 10000
n_plac e_f i e lds_pe r_c e l l :
   va lue s :
    − 1
opt imi z e r :
   va lue s :
− adam
place_c e l l_r f :
   va lue s :
    − 0. 09
    − 0. 12
    − 0. 15
    − 0. 18
    − 0. 21
    − 0. 24
place_f i e l d_l o s s :
   va lue s :
− c r o s s ent r opy
place_f i e ld_no rma l i z a t i on :
   va lue s :
− global
place_f i e ld_
   va lue s :
   va lue s :
− sof tmax_of_di f f e r enc e s
readout_dropout :
   va lue s :
    − 0
recur rent_dropout :
   va lue s :
    − 0
rnn_type :
   va lue s :
    − RNN
    − LSTM
    − UGRNN
    − GRU
seed:
   va lue s :
    − 0
    − 1
    − 2
sequence_length :
   va lue s :
    − 20
sur round_scale :
   va lue s :
    − 1. 5
    − 2
    − 2 . 5
    − 3
weight_decay :
   va lue s :
− 0.0001
~~~

### F.14 Multiple Scales and Multiple Fields Difference-of-Softmaxes Place

~~~
Cells
method : g r id
me t r i c :
  go al : minimize
  name : pos_decoding_err
parameter s :
Ng:
   v a lue s :
    − 1024
Np:
   v a lue s :
    − 512
a c t i v a t i o n :
   va lue s :
    − relu
batch_s ize :
   va lue s :
    − 200
bin_side_in_m :
   va lue s :
    − 0. 05
box_height_in_m:
   va lue s :
    − 2 . 2
box_width_in_m:
   va lue s :
    − 2 . 2
i n i t i a l i z e r :
49
   va lue s :
    − glorot_uniform
i s_periodic :
   va lue s :
    − f a l s e
l e a rning_r a t e :
   va lue s :
    − 0.0001
n_epochs :
   va lue s :
    − 20
n_grad_steps_per_epoch :
   va lue s :
    − 10000
n_plac e_f i e lds_pe r_c e l l :
   va lue s :
    − Poi s son ( 2 . 0)
    − Poi s son ( 3 . 0)
opt imi z e r :
   va lue s :
    − adam
place_c e l l_r f :
   va lue s :
    − 0. 12
    − Uniform( 0. 06, 0. 24 )
    − Uniform( 0. 06, 1. 0)
place_f i e l d_l o s s :
   va lue s :
    − c r o s s ent r opy
place_f i e ld_no rma l i z a t i on :
   va lue s :
− global
place_f i e ld_va lue s :
   va lue s :
− di f f e r enc e_o f_g aus s i ans
readout_dropout :
va lue s :
    − 0
recur rent_dropout :
   va lue s :
    − 0
rnn_type :
   va lue s :
    − RNN
    − UGRNN
seed:
   va lue s :
    − 0
    − 1
    − 2
sequence_length :
   va lue s :
    − 20
sur round_scale :
   va lue s :
    − 2
    − Uniform( 1. 50, 2. 50)
    − Uniform( 1. 25, 4. 50)
weight_decay :
   va lue s :
    − 0.0001− 0. 08
    − 0. 09
    − 0. 1
    − 0. 11
    − 0. 12
    − 0. 13
    − 0. 14
    − 0. 15
    − 0. 16
    − 0. 17
    − 0. 18
    − 0. 19
    − 0. 2
    − 0. 24
    − 0. 28
    − 0. 32
place_f i e l d_l o s s :
   va lue s :
    − c r o s s ent r opy
place_f i e ld_no rma l i z a t i on :
   va lue s :
    − global
place_f i e ld_va lue s :
   va lue s :
    − di f f e r enc e_o f_g aus s i ans
readout_dropout :
   va lue s :
    − 0
    − 0. 5
recur rent_dropout :
   va lue s :
    − 0
    − 0. 5
rnn_type :
   va lue s :
    − RNN
    − LSTM
    − UGRNN
    − GRU
seed:
   va lue s :
    − 0
    − 1
    − 2
sequence_length :
   va lue s :
    − 20
sur round_scale :
   va lue s :
    − 1. 5
    − 2
    − 2 . 5
    − 3
    − 4
weight_decay :
   va lue s :
    − 0.0001
~~~

### F.15 Nayebi et al. 2021[47] Replication

~~~
method : g r i d metric :
  goal : minimize
  name : pos_decoding_err
parameters :
Ng :
      values :
         – 4096
Np :
      values :
         – 512
activation :
      values :
         – relu
batch_size :
      values :
         – 200
bin_side_in_m :
      values :
         – 0. 05
box_height_in_m :
      values :
         – 2 . 2
box_width_in_m :
      values :
         – 2 . 2
initializer:
      values :
         – glorot_uniform i s _periodic :
      values :
         – false
learning _rate :
      values :
         – 0. 0001
n_epochs :
      values :
         – 20
n_grad_steps_per_epoch :
      values :
         – 10000
n_place_fields_per_cell :
      values :
         – 1
optimizer:
      values :
         – adam
place_cell _r f : \
      values :
         – 0. 12
place_f i e l d _l o s s :
      values :
         – crossentropy
place_f i e l d _n o r m a l i z a t i o n :
v alu e s :
         – global
place_f i e l d _v a l u e s :
      values :
         – difference_o f _g a u s s i a n s readout_dropout :
      values :
         – 0
recurrent_dropout :
      values :
         – 0
rnn_type :
      values :
         – RNN
         – LSTM
         – UGRNN
         – GRU
seed :
      values :
         – 0
         – 1
         – 2
         – 3
         – 4
sequence_length :
      values :
         – 20
surround_scale :
      values :
         – 2
weight_decay :
      values :
         – 0. 0001
~~~

## References

[1] Emergence of dynamically reconfigurable hippocampal responses by learning to perform proba-bilistic spatial reasoning. biorxiv.

[2] High-performing neural network models of visual cortex benefit from high latent dimensionality, July 2022. Pages: 2022.07.13.499969 Section: New Results.

[3] Andrea Banino, Caswell Barry, Benigno Uria, Charles Blundell, Timothy Lillicrap, Piotr Mirowski, Alexander Pritzel, Martin J. Chadwick, Thomas Degris, Joseph Modayil, Greg Wayne, Hubert Soyer, Fabio Viola, Brian Zhang, Ross Goroshin, Neil Rabinowitz, Razvan Pascanu, Charlie Beattie, Stig Petersen, Amir Sadik, Stephen Gaffney, Helen King, Koray Kavukcuoglu, Demis Hassabis, Raia Hadsell, and Dharshan Kumaran. Vector-based navigation using grid-like representations in artificial agents. Nature, 557(7705):429–433, May 2018.

[4] Pouya Bashivan, Kohitij Kar, and James J. DiCarlo. Neural population control via deep image synthesis. Science, 364(6439):eaav9436. May 2019. Publisher: American Association for the Advancement of Science.

[5] Romain Bourboulou, Geoffrey Marti, François-Xavier Michon, Elissa El Feghaly, Morgane Nouguier, David Robbe, Julie Koenig, and Jerome Epsztein. Dynamic control of hippocampal spatial coding resolution by local visual cues. Elife, 8:e44487, 2019.

[6] Tom B. Brown, Benjamin Mann, Nick Ryder, Melanie Subbiah, Jared Kaplan, Prafulla Dhariwal, Arvind Neelakantan, Pranav Shyam, Girish Sastry, Amanda Askell, Sandhini Agarwal, Ariel Herbert-Voss, Gretchen Krueger, Tom Henighan, Rewon Child, Aditya Ramesh, Daniel M. Ziegler, Jeffrey Wu, Clemens Winter, Christopher Hesse, Mark Chen, Eric Sigler, Mateusz Litwin, Scott Gray, Benjamin Chess, Jack Clark,Christopher Berner, Sam McCandlish, Alec Radford, Ilya Sutskever, and Dario Amodei. Language Models are Few-Shot Learners. arXiv:2005.14165 [cs], July 2020. arXiv: 2005.14165.

[7] Yoram Burak and Ila Fiete. Do We Understand the Emergent Dynamics of Grid Cell Activity? Journal of Neuroscience, 26(37):9352–9354, September 2006. Publisher: Society for Neuroscience Section: Journal Club.

[8] Yoram Burak and Ila R Fiete. Unpublished observations. 2008.

[9] Yoram Burak and Ila R. Fiete. Accurate Path Integration in Continuous Attractor Network Models of Grid Cells. PLOS Computational Biology, 5(2):e1000291, February 2009. Publisher: Public Library of Science.

[10] Yoram Burak and Ila R Fiete. Accurate path integration in continuous attractor network models of grid cells. PLoS Comput Biol, 5(2):e1000291, Feb 2009.

[11] Malcolm G. Campbell, Samuel A. Ocko, Caitlin S. Mallory, Isabel I. C. Low, Surya Ganguli, and Lisa M. Giocomo. Principles governing the integration of landmark and self-motion cues in entorhinal cortical codes for navigation. Nature Neuroscience, 21(8):1096–1106, August 2018. Number: 8 Publisher: Nature Publishing Group.

[12] Rishidev Chaudhuri, Berk Gerçek, Biraj Pandey, Adrien Peyrache, and Ila Fiete. The intrinsic attractor manifold and population dynamics of a canonical cognitive circuit across waking and sleep. Nat Neurosci, 22(9):1512–1520, 09 2019.

[13] Junyoung Chung, Caglar Gulcehre, KyungHyun Cho, and Yoshua Bengio. Empirical Evaluation of Gated Recurrent Neural Networks on Sequence Modeling. NIPS Workshop Deep Learning and Representation Learning, December 2014. Number: arXiv:1412.3555 arXiv:1412.3555 [cs].

[14] Jasmine Collins, Jascha Sohl-Dickstein, and David Sussillo. Capacity and Trainability in Recurrent Neural Networks. International Conference on Learning Representations, March 2017. Number: arXiv:1611.09913 arXiv:1611.09913 [cs, stat].

[15] Christopher J Cueva and Xue-Xin Wei. Emergence of grid-like representations by training recurrent neural networks to perform spatial localization. International Conference on Learning Representations, page 19, 2018.

[16] Nathan B Danielson, Jeffrey D Zaremba, Patrick Kaifosh, John Bowler, Max Ladow, and Attila Losonczy. Sublayer-specific coding dynamics during spatial navigation and learning in hippocampal area ca1. Neuron, 91(3):652–665, 2016.

[17] William Dorrell, Peter E Latham, Timothy EJ Behrens, and James CR Whittington. Actionable neural representations: Grid cells from minimal constraints. arXiv preprint arXiv:2209.15563, 2022.

[18] David Dupret, Joseph O’neill, Barty Pleydell-Bouverie, and Jozsef Csicsvari. The reorganization and reactivation of hippocampal maps predict spatial memory performance. Nature neuroscience, 13(8):995–1002, 2010.

[19] Tamir Eliav, Shir R. Maimon, Johnatan Aljadeff, Misha Tsodyks, Gily Ginosar, Liora Las, and Nachum Ulanovsky. Multiscale representation of very large environments in the hippocampus of flying bats. Science, 372(6545):eabg4020, May 2021. Publisher: American Association for the Advancement of Science.

[20] Jeffrey L. Elman. Finding Structure in Time. Cognitive Science, 14(2):179–211, 1990. _eprint: https://onlinelibrary.wiley.com/doi/pdf/10.1207/s15516709cog1402_1.

[21] Logan Engstrom, Andrew Ilyas, Shibani Santurkar, Dimitris Tsipras, Firdaus Janoos, Larry Rudolph, and Aleksander Madry. Implementation Matters in Deep Policy Gradients: A Case Study on PPO and TRPO. arXiv:2005.12729 [cs, stat], May 2020. arXiv: 2005.12729.

[22] Ila R Fiete, Yoram Burak, and Ted Brookings. What grid cells convey about rat location. J Neurosci, 28(27):6858–71, Jul 2008.

[23] Mark C Fuhs and David S Touretzky. A spin glass model of path integration in rat medial entorhinal cortex. J Neurosci, 26(16):4266–4276, 2006.

[24] Marianne Fyhn, Torkel Hafting, Alessandro Treves, May-Britt Moser, and Edvard I Moser. Hippocampal remapping and grid realignment in entorhinal cortex. Nature, 446(7132):190–194, 2007.

[25] Richard J. Gardner, Erik Hermansen, Marius Pachitariu, Yoram Burak, Nils A. Baas, Benjamin A. Dunn, May-Britt Moser, and Edvard I. Moser. Toroidal topology of population activity in grid cells. Nature, 602(7895):123–128, February 2022. Number: 7895 Publisher: Nature Publishing Group.

[26] Richard J. Gardner, Li Lu, Tanja Wernle, May-Britt Moser, and Edvard I. Moser. Correlation structure of grid cells is preserved during sleep. Nature Neuroscience, 22(4):598–608, April 2019. Number: 4 Publisher: Nature Publishing Group.

[27] Jeffrey L Gauthier and David W Tank. A dedicated population for reward coding in the hippocampus. Neuron, 99(1):179–193, 2018.

[28] Joshua I. Glaser, Ari S. Benjamin, Raeed H. Chowdhury, Matthew G. Perich, Lee E. Miller, and Konrad P. Kording. Machine Learning for Neural Decoding. eNeuro, 7(4), July 2020. Publisher: Society for Neuroscience Section: Research Article: Methods/New Tools.

[29] Yi Gu, Sam Lewallen, Amina A Kinkhabwala, Cristina Domnisoru, Kijung Yoon, Jeffrey L Gauthier, Ila R Fiete, and David W Tank. A map-like micro-organization of grid cells in the medial entorhinal cortex. Cell, 175(3):736–750.e30, 10 2018.

[30] Torkel Hafting, Marianne Fyhn, Sturla Molden, May-Britt Moser, and Edvard I. Moser. Microstructure of a spatial map in the entorhinal cortex. Nature, 436(7052):801–806, August 2005. Number: 7052 Publisher: Nature Publishing Group.

[31] Phil A Hetherington and Matthew L Shapiro. Hippocampal place fields are altered by the removal of single visual cues in a distance-dependent manner. Behavioral neuroscience, 111(1):20, 1997.

[32] Geoffrey Hinton, Nitish Srivastava, and Kevin Swersky. Lecture 6e-RMSProp.

[33] Sepp Hochreiter and Jürgen Schmidhuber. Long Short-Term Memory. Neural Computation, 9(8):1735–1780, November 1997.

[34] Stig A Hollup, Sturla Molden, James G Donnett, May-Britt Moser, and Edvard I Moser. Accumulation of hippocampal place fields at the goal location in an annular watermaze task. Journal of Neuroscience, 21(5):1635–1644, 2001.

[35] Andrew Ilyas, Logan Engstrom, Shibani Santurkar, Dimitris Tsipras, Firdaus Janoos, Larry Rudolph, and Aleksander Madry. A Closer Look at Deep Policy Gradients. arXiv:1811.02553 [cs, stat], May 2020. arXiv: 1811.02553.

[36] John Jumper, Richard Evans, Alexander Pritzel, Tim Green, Michael Figurnov, Olaf Ronneberger, Kathryn Tunyasuvunakool, Russ Bates, Augustin Žídek, Anna Potapenko, Alex Bridgland, Clemens Meyer, Simon A. A. Kohl, Andrew J. Ballard, Andrew Cowie, Bernardino Romera-Paredes, Stanislav Nikolov, Rishub Jain, Jonas Adler, Trevor Back, Stig Petersen, David Reiman, Ellen Clancy, Michal Zielinski, Martin Steinegger, Michalina Pacholska, Tamas Berghammer, Sebastian Bodenstein, David Silver, Oriol Vinyals, Andrew W. Senior, Koray Kavukcuoglu, Pushmeet Kohli, and Demis Hassabis. Highly accurate protein structure prediction with AlphaFold. Nature, 596(7873):583–589, August 2021. Number: 7873 Publisher: Nature Publishing Group.

[37] I. Kanitscheider and I. R. Fiete. Training recurrent networks to generate hypotheses about how the brain solves hard navigation problems. Advances in Neural Information Processing Systems (NeurIPS), 2017.

[38] Mikail Khona, Sarthak Chandra, and Ila R. Fiete. From smooth cortical gradients to discrete modules: spontaneous and topologically robust emergence of modularity in grid cells. bioRxiv, page 2021.10.28.466284, January 2022.

[39] Mikail Khona and Ila R Fiete. Attractor and integrator networks in the brain. preprint at https://arxiv.org/abs/2112.03978, 2021.

[40] Timothy D. Kim, Thomas Z. Luo, Jonathan W. Pillow, and Carlos D. Brody. Inferring Latent Dynamics Underlying Neural Population Activity via Neural Differential Equations. In Proceedings of the 38th International Conference on Machine Learning, pages 5551–5561. PMLR, July 2021. ISSN: 2640-3498.

[41] Diederik P. Kingma and Jimmy Ba. Adam: A Method for Stochastic Optimization. International Conference on Learning Representations, January 2017. Number: arXiv:1412.6980 arXiv:1412.6980 [cs].

[42] Ashok Litwin-Kumar, Kameron Decker Harris, Richard Axel, Haim Sompolinsky, and L. F. Abbott. Optimal Degrees of Synaptic Connectivity. Neuron, 93(5):1153–1164.e7, March 2017.

[43] Jesse A. Livezey, Kristofer E. Bouchard, and Edward F. Chang. Deep learning as a tool for neural data analysis: Speech classification and cross-frequency coupling in human sensorimotor cortex. PLOS Computational Biology, 15(9):e1007091, September 2019. Publisher: Public Library of Science.

[44] A. Mathis, A. Herz, and M. Stemmler. Optimal population codes for space: grid cells outperform place cells. Neural Comp., 24:2280–2317, 2012.

[45] Alexander Mathis, Andreas V. M. Herz, and Martin Stemmler. Optimal Population Codes for Space: Grid Cells Outperform Place Cells. Neural Computation, 24(9):2280–2317, September 2012.

[46] Alexander Mathis, Pranav Mamidanna, Kevin M. Cury, Taiga Abe, Venkatesh N. Murthy, Mackenzie Weygandt Mathis, and Matthias Bethge. DeepLabCut: markerless pose estimation of user-defined body parts with deep learning. Nature Neuroscience, 21(9):1281–1289, September 2018. Number: 9 Publisher: Nature Publishing Group.

[47] Aran Nayebi, Alexander Attinger, Malcolm Campbell, Kiah Hardcastle, Isabel Low, Caitlin S Mallory, Gabriel Mel, Ben Sorscher, Alex H Williams, Surya Ganguli, Lisa Giocomo, and Dan Yamins. Explaining heterogeneity in medial entorhinal cortex with task-driven neural networks. In Advances in Neural Information Processing Systems, volume 34, pages 12167–12179. Curran Associates, Inc., 2021.

[48] J. O’Keefe and J. Dostrovsky. The hippocampus as a spatial map: Preliminary evidence from unit activity in the freely-moving rat. Brain Research, 34:171–175, 1971. Place: Netherlands Publisher: Elsevier Science.

[49] John O’Keefe and Dulcie H Conway. Hippocampal place units in the freely moving rat: why they fire where they fire. Experimental brain research, 31(4):573–590, 1978.

[50] Talmo D. Pereira, Diego E. Aldarondo, Lindsay Willmore, Mikhail Kislin, Samuel S.-H. Wang, Mala Murthy, and Joshua W. Shaevitz. Fast animal pose estimation using deep neural networks. Nature Methods, 16(1):117–125, January 2019. Number: 1 Publisher: Nature Publishing Group.

[51] P. D. Rich, H.-P. Liaw, and A. K. Lee. Large environments reveal the statistical structure governing hippocampal representations. Science, 345(6198):814–817, August 2014.

[52] Muhammad Saif-ur Rehman, Robin Lienkämper, Yaroslav Parpaley, Jörg Wellmer, Charles Liu, Brian Lee, Spencer Kellis, Richard Andersen, Ioannis Iossifidis, Tobias Glasmachers, and Christian Klaes. SpikeDeeptector: a deep-learning based method for detection of neural spiking activity. Journal of Neural Engineering, 16(5):056003, July 2019. Publisher: IOP Publishing.

[53] Masaaki Sato, Kotaro Mizuta, Tanvir Islam, Masako Kawano, Yukiko Sekine, Takashi Takekawa, Daniel Gomez-Dominguez, Alexander Schmidt, Fred Wolf, Karam Kim, et al. Distinct mechanisms of over-representation of landmarks and rewards in the hippocampus. Cell reports, 32(1):107864, 2020.

[54] Andrew Saxe, Stephanie Nelli, and Christopher Summerfield. If deep learning is the answer, what is the question? Nature Reviews Neuroscience, 22(1):55–67, January 2021. Number: 1 Publisher: Nature Publishing Group.

[55] Rylan Schaeffer, Mikail Khona, Leenoy Meshulam, Brain Laboratory International, and Ila Fiete. Reverse-engineering recurrent neural network solutions to a hierarchical inference task for mice. Advances in Neural Information Processing Systems, 33:4584–4596, 2020.

[56] Vemund Sigmundson Schøyen, Markus Borud Pettersen, Konstantin Holzhausen, Anders Malthe-Sørensen, and Mikkel Elle Lepperød. Navigating multiple environments with emergent grid cell remapping. bioRxiv, 2022.

[57] David Silver, Aja Huang, Chris J. Maddison, Arthur Guez, Laurent Sifre, George van den Driessche, Julian Schrittwieser, Ioannis Antonoglou, Veda Panneershelvam, Marc Lanctot, Sander Dieleman, Dominik Grewe, John Nham, Nal Kalchbrenner, Ilya Sutskever, Timothy Lillicrap, Madeleine Leach, Koray Kavukcuoglu, Thore Graepel, and Demis Hassabis. Mastering the game of Go with deep neural networks and tree search. Nature, 529(7587):484–489, January 2016. Number: 7587 Publisher: Nature Publishing Group.

[58] Trygve Solstad, Charlotte N. Boccara, Emilio Kropff, May-Britt Moser, and Edvard I. Moser. Representation of Geometric Borders in the Entorhinal Cortex. Science, 322(5909):1865–1868, December 2008. Publisher: American Association for the Advancement of Science.

[59] Ben Sorscher, Gabriel C Mel, Surya Ganguli, and Samuel A Ocko. A unified theory for the origin of grid cells through the lens of pattern formation. Advances in Neural Information Processing Systems, page 18, 2019.

[60] Ben Sorscher, Gabriel C. Mel, Samuel A. Ocko, Lisa Giocomo, and Surya Ganguli. A unified theory for the computational and mechanistic origins of grid cells. Technical report, bioRxiv, December 2020. Section: New Results Type: article.

[61] Sameet Sreenivasan and Ila Fiete. Grid cells generate an analog error-correcting code for singularly precise neural computation. Nat Neurosci, 14(10):1330–7, Sep 2011.

[62] Hanne Stensola, Tor Stensola, Trygve Solstad, Kristian Frøland, May-Britt Moser, and Edvard I. Moser. The entorhinal grid map is discretized. Nature, 492(7427):72–78, December 2012. Number: 7427 Publisher: Nature Publishing Group.

[63] David Sussillo and Omri Barak. Opening the Black Box: Low-Dimensional Dynamics in High-Dimensional Recurrent Neural Networks. Neural Computation, 25(3):626–649, March 2013.

[64] S.G. Trettel, J.B. Trimper, E. Hwaun, I.R. Fiete, and L.L. Colgin. Grid cell co-activity patterns during sleep reflect spatial overlap of grid fields during active behaviors. Nat Neurosci, 22(4):609–617, 04 2019.

[65] George Tucker, Surya Bhupatiraju, Shixiang Gu, Richard E. Turner, Zoubin Ghahramani, and Sergey Levine. The Mirage of Action-Dependent Baselines in Reinforcement Learning. arXiv:1802.10031 [cs, stat], November 2018. arXiv: 1802.10031.

[66] Greta Tuckute, Jenelle Feather, Dana Boebinger, and Josh H McDermott. Many but not all deep neural network audio models capture brain responses and exhibit hierarchical region correspondence. bioRxiv, 2022.

[67] Jakob Voigts, Ingmar Kanitscheider, Nicholas J. Miller, Enrique H. S. Toloza, Jonathan P. Newman, Ila R. Fiete, and Mark T. Harnett. Spatial reasoning via recurrent neural dynamics in mouse retrosplenial cortex. biorxiv, April 2022.

[68] James C. R. Whittington, Timothy H. Muller, Shirley Mark, Guifen Chen, Caswell Barry, Neil Burgess, and Timothy E. J. Behrens. The Tolman-Eichenbaum Machine: Unifying Space and Relational Memory through Generalization in the Hippocampal Formation. Cell, 183(5):1249–1263.e23, November 2020.

[69] SI Wiener, CA Paul, and H Eichenbaum. Spatial and behavioral correlates of hippocampal neuronal activity. Journal of Neuroscience, 9(8):2737–2763, 1989.

[70] Dehong Xu, Ruiqi Gao, Wen-Hao Zhang, Xue-Xin Wei, and Ying Nian Wu. Conformal isometry of lie group representation in recurrent network of grid cells. arXiv preprint arXiv:2210.02684, 2022.

[71] Daniel LK Yamins, Ha Hong, Charles F Cadieu, Ethan A Solomon, Darren Seibert, and James J DiCarlo. Performance-optimized hierarchical models predict neural responses in higher visual cortex. Proceedings of the national academy of sciences, 111(23):8619–8624, 2014.

[72] K.J. Yoon, M.A. Buice, R. Barry, C.and Hayman, N. Burgess, and I.R. Fiete. Specific evidence of low-dimensional continuous attractor dynamics in grid cells. Nat Neurosci, 16(8):1077–84, Aug 2013.

[73] Jeffrey D Zaremba, Anastasia Diamantopoulou, Nathan B Danielson, Andres D Grosmark, Patrick W Kaifosh, John C Bowler, Zhenrui Liao, Fraser T Sparks, Joseph A Gogos, and Attila Losonczy. Impaired hippocampal place cell dynamics in a mouse model of the 22q11. 2 deletion. Nature neuroscience, 20(11):1612–1623, 2017.

